# Structures and broad-spectrum growth-inhibiting activity of formomarinobactin, formylated marinobactin analogues from the *Pseudomonas lutea* clade

**DOI:** 10.64898/2026.01.14.699459

**Authors:** C. Grosse, K. Hughes, M. Lavender, B. Cornu, N. Brandt, S. Matthijs

## Abstract

*Pseudomonas graminis* LMG 21661^T^, an environmental strain of the *P. lutea* clade, produces the siderophore formomarinobactin, a novel marinobactin-like siderophore. Mass spectrometry revealed that formomarinobactin shares the same six-residue peptide backbone as marinobactin but contains formylated rather than acetylated N-hydroxyornithines. Alike marinobactins, formomarinobactins are produced as a suite of siderophores with a conserved hexapeptide core but varying lipid tail lengths (C10-C14), shorter than the C12 to C18 characteristic of marinobactins. Both the biosynthesis and cognate receptor genes of the formomarinobactin system in *P. graminis* are iron regulated but unaffected by zinc or nickel underscoring their role in iron homeostasis.

Genome mining combined with mass analyses demonstrated that formomarinobactin production is a conserved trait across the *P. lutea* clade, with one exception which appears to represent an intraspecific cheater that has lost siderophore production. Beyond the producing strains themselves, we identified a widespread distribution of putative formomarinobactin receptors among diverse *Pseudomonas* species, revealing a substantial capacity within the *P. fluorescens* super clade to pirate formomarinobactin as an iron source. Putative receptors were also found in genera outside the *Pseudomonas* genus. Growth stimulation assays confirmed functional formomarinobactin uptake in several *Pseudomonas* spp. and a *Phytopseudomonas* strain, with genetic validation in *Pseudomonas rhodesiae*.

Importantly, formomarinobactin production confers more than a nutritional advantage. Members of the *P. lutea* clade producing formomarinobactin display pronounced growth inhibiting activity against a broad spectrum of clinical and environmental Gram-positive and with lower efficacy against Gram-negative bacteria. Purified formomarinobactin was able to inhibit growth under iron-limiting conditions and to a lesser extent in iron-rich conditions, highlighting a dual role for this molecule in both iron acquisition and microbial growth inhibition.

## 1 Introduction

*Pseudomonas* constitutes a clade of ubiquitous Gram-negative bacteria renowned for their metabolic versatility and capability to colonize a wide range of environments (Silby et al. 2011). Species have been isolated from terrestrial (plants, insects, soil, fruits, human tissues) and marine (both seawater and freshwater) environments (Butaitė et al. 2017; Girard et al. 2021; 2023; Mulet et al. 2021; Silby et al. 2011). The genus increased exponentially with the description of 70 novel species in 10 years and over 400 species are validly published in the ICNP to date, excluding synonyms and misspellings (Freese et al., 2025, accessed November 20^th^, 2025).

To adapt to different natural environments in iron-limited conditions, *Pseudomonas* notably rely on specialized metabolites known as siderophores (Cornelis 2010). Indeed, iron availability is scarce due to low solubility of natural Fe^3+^ under aerobic conditions and neutral pH (Andrews et al. 2003). The most extensively studied siderophores of the genus are pyoverdines, high iron affinity molecules produced by fluorescent *Pseudomonas* species that play a key role in iron acquisition as well as virulence, and microbial competition (Cornelis et al. 2023; Meyer et al. 1996; Meyer 2000). These Non-Ribosomal Peptide Synthetases (NRPS)-regulated molecules chelate ferric iron (Fe³⁺) with strong specificity and are transported into the cell via dedicated outer-membrane receptors, enabling bacteria to thrive in iron-limited environments (Cornelis 2010; Ghssein and Ezzeddine 2022; Ravel and Cornelis 2003).

However, non-fluorescent *Pseudomonas* species have also developed iron chelating strategies by producing alternative, usually lower-affinity, siderophores (Cornelis and Matthijs, 2002). Although these systems have been less extensively characterized, they also play significant roles in ecological fitness, metal homeostasis, and interactions with plants and other microbes.

*P. graminis,* a member of the non-fluorescent *P. lutea* clade, was initially isolated from the phyllosphere of grasses (Behrendt et al. 1999; Peix et al. 2004) and later from apples (Alegre et al., 2013a) and cloud water (Besaury et al. 2017; Wirgot et al. 2019). Strains CPA-7 and 49M have been consistently studied for their use in the preservation of fresh cut fruits, i.e. apples, pears and melon, or *in planta*, where they have shown varying growth inhibition activity against *Escherichia coli* O157:H7, *Salmonella* Michigan, *Listeria innocua* and *Erwinia amylovora* (Abadias et al., 2014; Alegre et al., 2013a; Alegre et al., 2013b; Collazo et al., 2018; Iglesias et al., 2018; Mikiciński et al., 2016). Investigating the mechanisms involved in their antagonism raised hypotheses concerning the involvement of siderophores (Mikiciński et al. 2016) or the reliance on mere carbon competition (Collazo et al. 2017). Further genome sequencing of strain UASWS1507 isolated from an apple tree in Switzerland (Crovadore et al. 2016), PDD-13b-3 from cloud water in France (Besaury et al. 2017), and CPA-7 from apples (Collazo et al. 2025), and analysis of the latter two genomes, highlighted the presence of a siderophore cluster which was not successfully annotated or elucidated by the authors. Nonetheless, Besaury et al., suggested toward pyoverdine production, however *P. graminis* is non-fluorescent so likely produces a non-fluorescent type siderophore (Meyer et al., 2002).

The aim of our research was to characterize, both in structure and biological activity, iron-regulated molecules produced by *Pseudomonas graminis*. We were able to identify a NRPS biosynthetic gene cluster *in silico* and determine the structure of the suite of siderophores produced by LC-HRMS. Our comparative genome and LC-MS analyses with other strains of the *P. lutea* clade indicated its presence in these strains, associated with a functional production and a dedicated receptor. The suite of siderophores produced was shown to correspond to novel lipopeptides differing in lipid tail length, named formomarinobactins. They exhibited both iron chelating and growth inhibiting activities against Gram-positive and to a lesser Gram-negative environmental and clinical strains. These findings suggest that *P. graminis* and members of the *P. lutea* clade though non-fluorescent, produce a high affinity siderophore involved in their colonization of environmental habitats.

## 2 Material and Methods

### 2.1 Strains and growth conditions

All the strains used in this study are reported in Table 1. *Pseudomonas* strains were cultured in iron-poor Casamino Acids medium (CAA) at 28°C. The medium was prepared with 5 g/L Bacto Casamino Acids (Difco Laboratories), 0.9 g/L K_2_HPO_4_, 0.25 g/L MgSO_4_.7H_2_O. Target organism strains were cultured in a rich (not iron depleted) in-house medium 856 at 37°C (Matthijs et al. 2013) for the *in vitro* antagonism assays. To obtain solid medium for both CAA and 856, agar was added at a final concentration of 15 g/L. For the soft agar overlay, 856 medium was supplemented with agar at 7.5 g/L. For 2,2’-bipyridine plates, 2,2’-bipyridine was directly added to melted CAA cooled down to 60°C at optimal concentrations according to each strain studied.

**Table 1.**
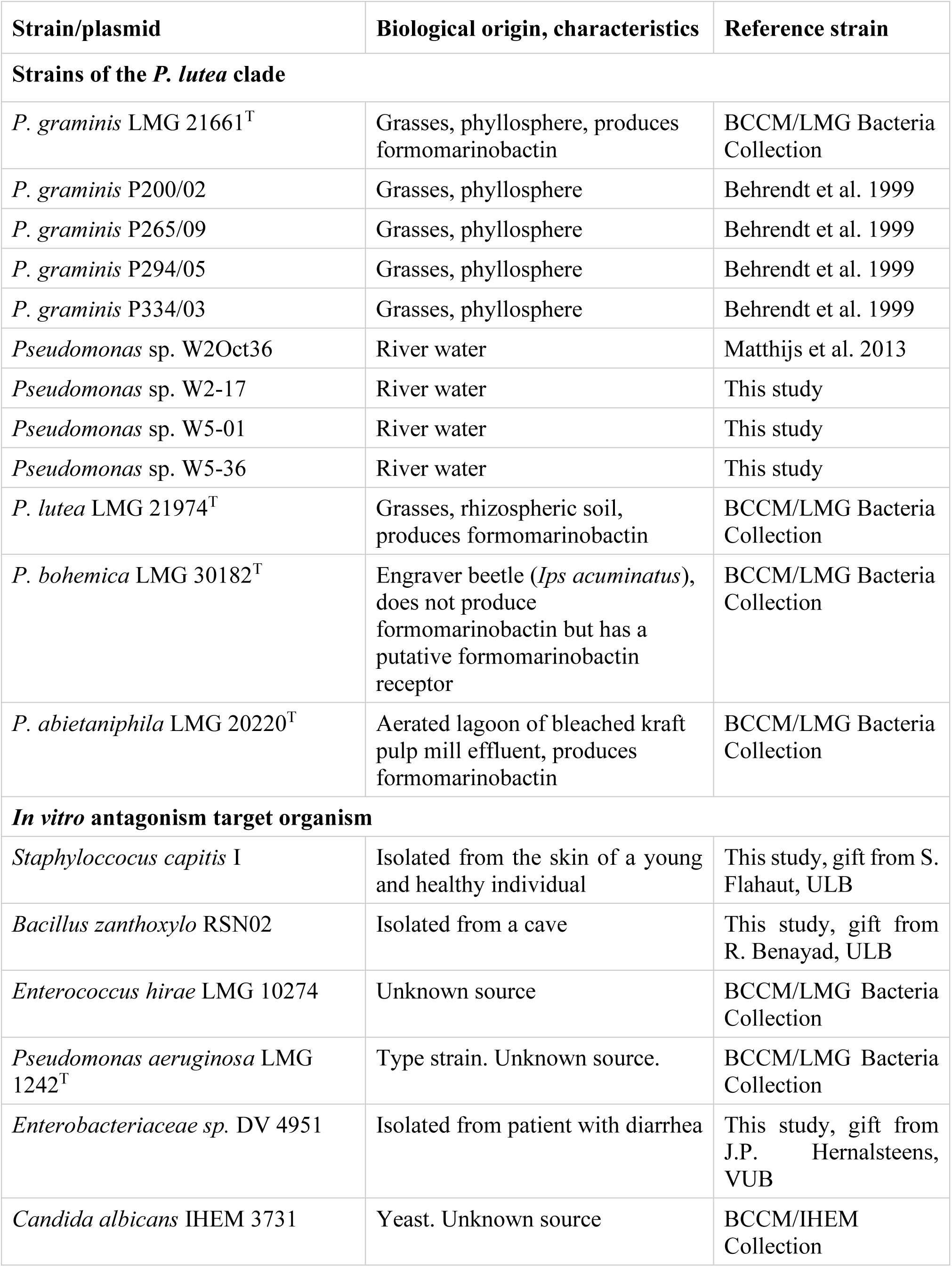
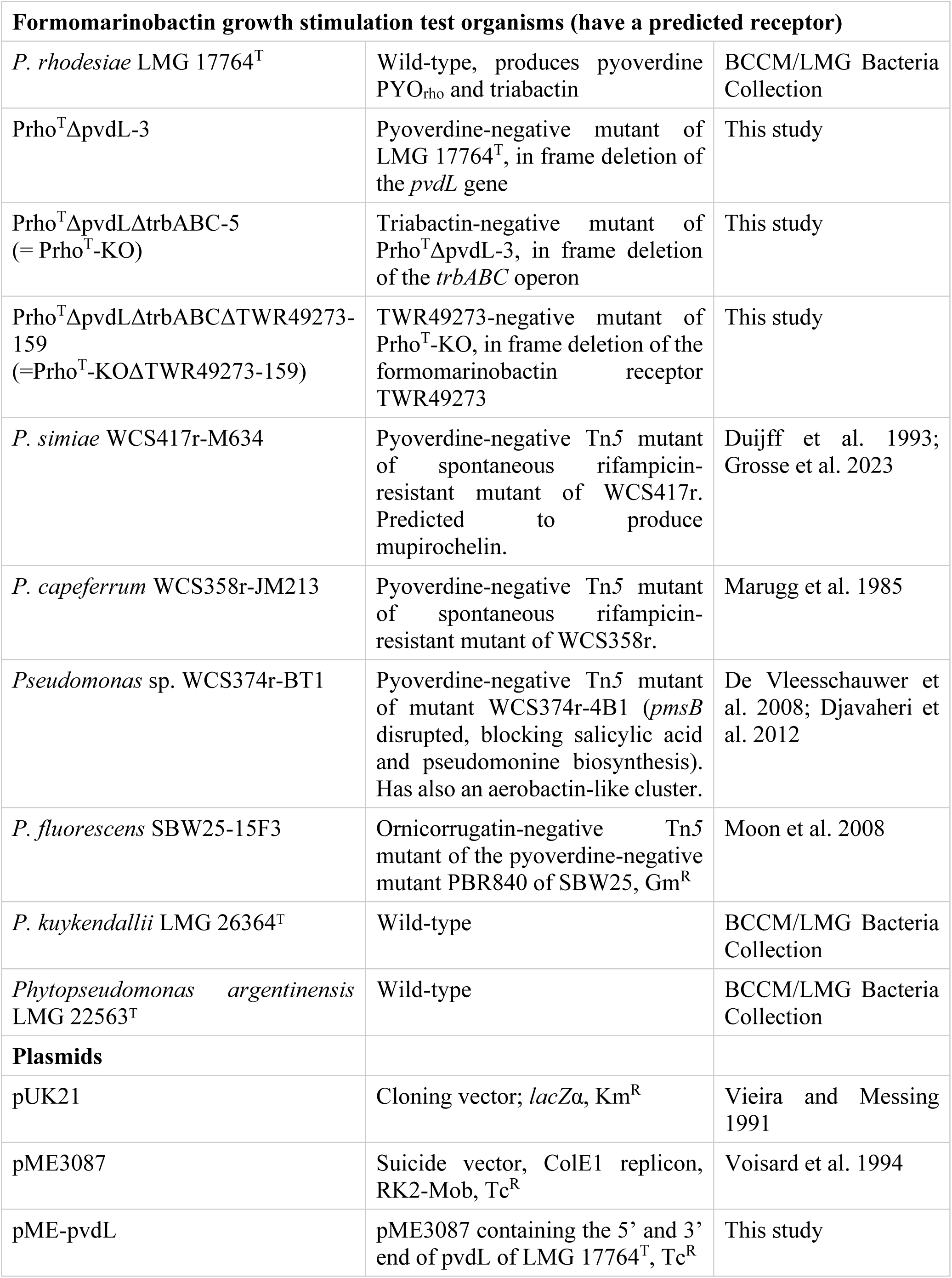
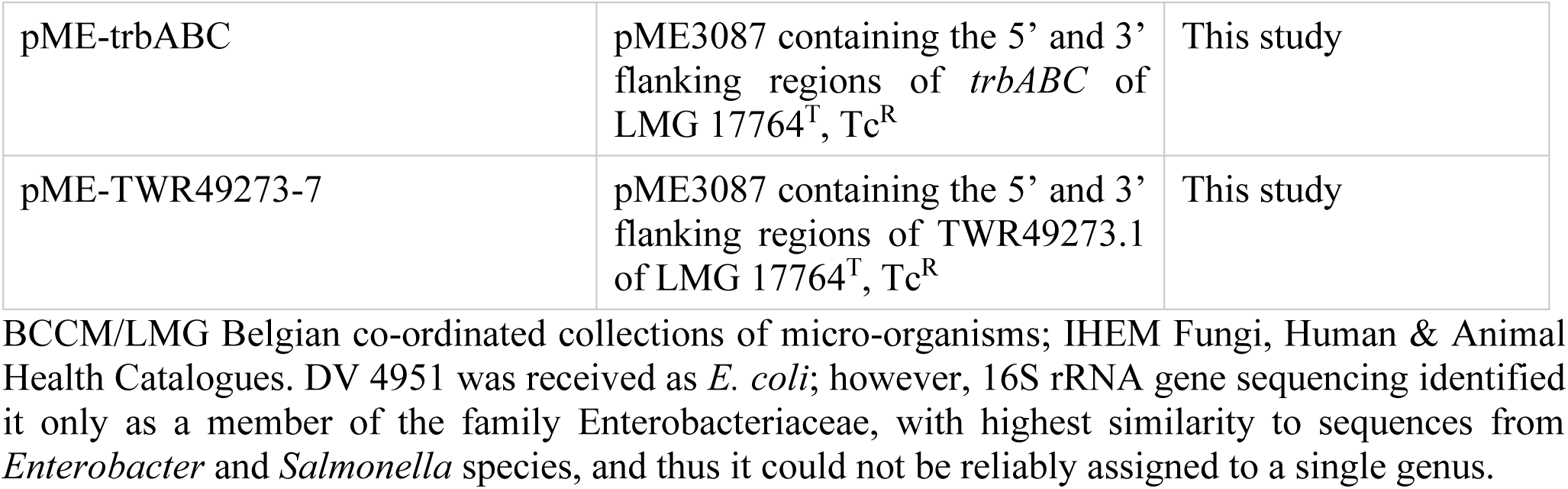
List of strains and plasmids used in this study.

### 2.2 *In silico* analysis of *P. graminis* LMG 21661^T^ siderophore gene cluster

Siderophore gene cluster analyses were conducted using BLAST (https://www.ncbi.nlm.nih.gov), PKS/NRPS analysis (http://nrps.igs.umaryland.edu), Interpro (https://www.ebi.ac.uk/interpro) and the NRPS/PKS module analysis feature implemented in antiSMASH (https://antismash.secondarymetabolites.org).

### 2.3 RT- qPCR assay of *P. graminis* LMG 21661^T^ grown in different iron conditions

RT-qPCR analyses were performed to determine whether the gene cluster identified *in silico* was expressed and regulated by iron. To do so, *P. graminis* LMG 21661^T^ was grown overnight in liquid CAA medium and 20 mL of the preculture, adjusted to OD_660nm_ = 0.2, was inoculated in 180 mL of liquid CAA and incubated at 30°C under agitation at 200 rpm, till the OD_660nm_ reached 0.2. The culture was divided into six subcultures to study the effect of increasing iron concentrations. Consequently, the subcultures were supplemented with FeCl_3_ (Merck), ZnCl_2_ (Fisher Chemical) or NiCl_2_ (Fluka) to a final concentration of 0, 0.5, 1, 2, 4 and 8 µM. A refined concentration gradient of 0, 0.05, 0.1, 0.2, 0.5 and 0.7 μM FeCl_3_ was tested. These supplemented subcultures were then incubated for 1h at 28°C (200 rpm). Thereafter, the subcultures were centrifuged, resuspended in fresh liquid CAA and RNAprotect Bacteria reagent (Qiagen) was added. The total RNA was extracted with a RNeasy Mini kit (Qiagen). The samples were treated with RNAse free DNA Set (Qiagen) to eliminate presumptive genomic DNA contamination. The integrity and purity of RNA were assessed by electrophoresis on a 1% Agarose gel and using a NanoDrop DeNovix DS-11 spectrophotometer (DeNovix Inc., Wilmington, DE USA). Reverse transcription was done with 1 µg of RNA for each sample with RevertAid H Minus First Strand cDNA synthesis kit with oligo(dt) primers (Thermo Scientific). The cDNA was purified by means of the QIAquick PCR Purification kit (Qiagen). Quantification of the cDNA was conducted using Fast start Essential DNA Green Master (Roche Applied Science) and the appropriate primers (Table 2) on a LightCycler 96 (Roche). The studied genes were *fmbD*, the first NRPS gene, and *fmbG*, the receptor gene. The quantity of DNA was normalized against the expression of *nadB* (L-aspartate oxidase), chosen as reference gene. Genomic DNA was extracted from LMG 21661^T^ using the DNeasy Blood and Tissue Kit (Qiagen) of a culture grown overnight in 856 media. A standard curve was generated using five different dilutions of genomic DNA and was used to quantify samples. RT-qPCR analyses were conducted at least in triplicate.

**Table 2.**
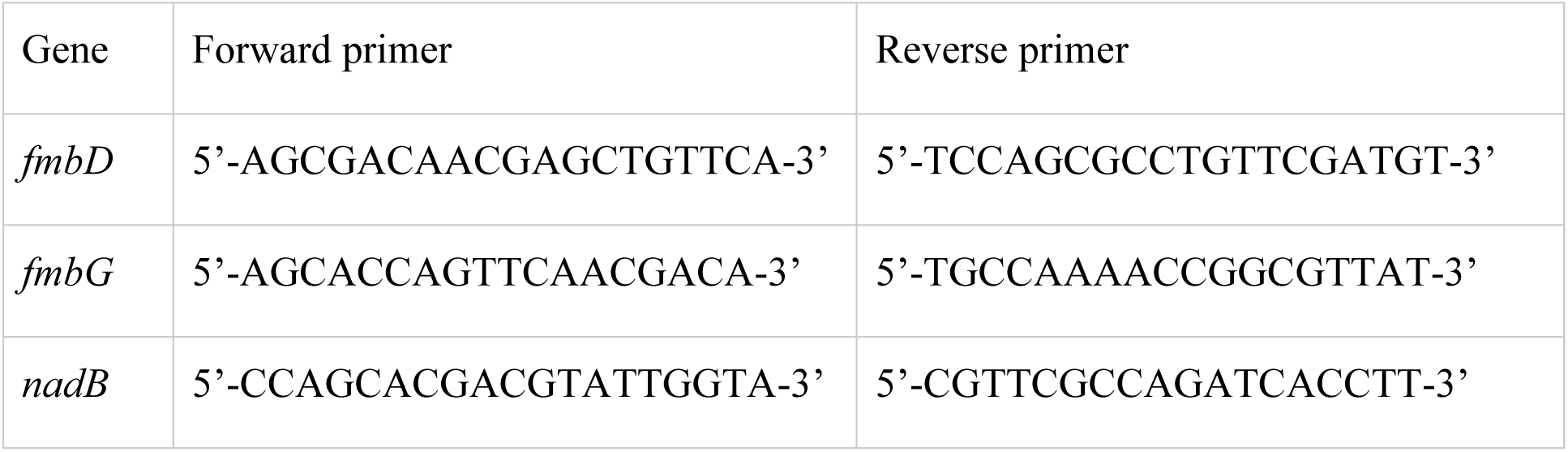
Sequence of the primers used for RT-qPCR.

### 2.4 LC-MS analysis of a siderophore suite produced by *P. graminis* LMG 21661^T^

Siderophore production was evaluated by CAS assay (Schwyn and Neilands 1987). *P. graminis* LMG 21661^T^ was analyzed by LC-MS to determine the nature of the siderophore. The strain was grown for 48 h at 28^°^C in 100 mL of CAA. Subsequently, the cultures were centrifuged for 15 min at 10,000g (Sorvall Lynx 6000, Thermoscientific). The filtered supernatant (0.2 µM, Millisart Satorius filter, Stedim Biotech) was applied to a CHROMABOND C18 ecf column (Macherey-Nagel). Elution of the compounds of interest was performed with acetonitrile/H_2_O (70:30 v/v). The purified fractions were analyzed by LC-MS using an Alliance e2695 separation module (Waters) coupled to an Altus SQ mass detector (Perkin Elmer). The method used consisted of solvents A and B, water and acetonitrile, respectively, both containing 0.1% formic acid. Separation was achieved on a C18 Altima column (250 x 4.6 mm, 10 μm, Grace), at a flow rate of 0.5 mL/ min, with a gradient of 10% B to 70% B in 30 min. This was followed by equilibration for 5 min at 10% B. The samples were analyzed by ESI in positive mode. The MS parameters were cone voltage 49 V, capillary voltage 3.1 kV, source temperature 150°C, gas desolvation temperature 600°C. Full scan *m/z* was acquired between 400 to 1800 with a scan rate of 1200 Da/sec.

### 2.5 Large scale purification of siderophores produced by *P. graminis* LMG 21661^T^

The siderophore variants observed by LC-MS were further purified on a large scale. Thirty milliliters (10 x 3 mL) of a *P. graminis* LMG 21661^T^ overnight preculture grown in 100 mL CAA were used to inoculate 10L (10 x 1L) of Minimal Succinate Medium (Vindeirinho et al. 2021). The flasks were then incubated for 72h at 28^°^C under agitation (180 rpm). The cultures were centrifuged at 4500 rpm for 30 min (Sorvall RC 12BP Plus, Thermo Scientific). The supernatants were pooled and passed through a CHROMABOND C18 ecf column (Macherey-Nagel) (2.5 x 4 cm). Salts were removed by washing the column with 200 mL of mQ water. The siderophores were then eluted with 100 mL of 70% acetonitrile. The fraction obtained was filtered (0.2 µM, Millisart), evaporated then lyophilized. The purification step was repeated 4 times. The semi-purified fraction was further purified using a SunFire Prep C18 column (19 x 250 mm, 5 µM, Waters) connected to an LC Prep 150 system (Waters) equipped with a UV/Vis 2489 detector (Waters). The solvents used were A, water, and solvent B, methanol. The method was a gradient set at a flow rate of 12 mL/min from 40% B to 90% B in 12 min, followed by 4 min at 100% B then column re-equilibration at 40% B for 5 min. UV detection was performed at 220 nm. Automated fraction collection using a Waters Fraction Collector III Collection was set at 80% tube volume between 3 and 15 min. Fractions were analyzed by LC-MS according to an adapted version of the method described in section (LC-MS analysis of siderophores) and pooled together according to similarities in chromatogram profiles (same *m/z* ion and retention time).

### 2.6 Structural determination of purified siderophore of *P. graminis* LMG 21661^T^ by LC-ESI-qTOF

The structure of the purified siderophores was further investigated using an LC 1200 (Agilent) chain coupled to a QTOF-6520 (Agilent). Chromatographic separation was conducted on an C18 Altima Grace Column (250 x 4.6 mm, 10 μm, Grace) maintained at 30°C. Solvents A and B consisted of water and acetonitrile, respectively, with 0.1% formic acid for both and set to a flow rate of 0.5 mL/min. The method started at 30%B held for 2 min, followed by a gradient of 30% B to 90%B in 25 min, 100%B was passed through the column for 4 min, then the column was brought back to initial conditions at 30%B in 1 minute and maintained for 5 min. The full method duration was 37 min. The analysis was performed in positive ion mode with electrospray inlet parameters set to 325°C, 7 L/min and 45 psig for gas temperature, drying gas flow and nebulizer, respectively. Capillary voltage was set to 4.5 kV, fragmentor to 175 V and skimmer to 65 V. Full scan HRMS spectra were recorded between *m*/*z* 100 and 1000 and MS/MS from *m/z* 50 to 1000. Targeted MS/MS fragmentation was performed by CID at 55 V for *m/z* 874.4177, 876.4379, 902.4542, 904.4655 and at 65 V for *m/z* 930.4878 and 932.5002, with an isolation window of 4 *m/z*. A second untargeted MS/MS analysis was conducted at 35 eV. MS raw data were converted to .mzML using MSConvert (ProteoWizard) and processed using MZMine 4.

### 2.7 Creation of a Custom Reference Genome Dataset for the *Pseudomonas lutea* clade

Genome sequence data for *Pseudomonas* strains belonging to the *P. lutea* clade were retrieved from the NCBI RefSeq database (query date: 5 December 2025) (Sayers et al. 2024). Genome quality was evaluated using CheckM v1.2.5 (Parks et al. 2015), retaining only genomes with ≥95% completeness and ≤5% contamination. In total, 46 genomes were selected, comprising 6 complete and 40 draft assemblies, and imported into a custom ‘PluteaC’ database using BioNumerics v7.6.3 (Applied Maths, bioMérieux).

### 2.8 Genome sequencing of river isolates

The genomes of *Pseudomonas* sp. W2-17, *Pseudomonas* sp. W5-01, *Pseudomonas* sp. W5-36 and *Pseudomonas* sp. W2Oct36 were sequenced. Therefore, genomic DNA was isolated using a Maxwell® RSC Cultured Cells DNA Kit (cat.# AS1620) and a Maxwell® RSC Instrument (cat.# AS1620), after a prior enzymatic lysis step. The integrity and purity of the DNA was evaluated on a 1.0% (w/v) agarose gel and by spectrophotometric measurements at 234, 260 and 280 nm. A Quantus™ fluorimeter and a QuantiFluor™ ONE dsDNA System (Promega Corporation, Madison, WI, USA) were used to estimate the DNA concentration. Library preparation and genome sequencing were performed by SeqCoast Genomics (Portsmouth, USA). Library preparation was performed using an Illumina DNA Prep tagmentation kit and unique dual indexes. Paired-end sequence reads (2×150bp) were generated using a NextSeq 2000 platform (Illumina Inc., San Diego, CA, USA). Quality check, trimming of the raw sequence reads and genome assembly (*de novo*) were performed using the Shovill v1.1.0 pipeline (https://github.com/tseemann/shovill), which uses SPAdes v3.15.4 (Bankevich et al. 2012) as its core and aims at a sequencing depth of 150x. Contigs shorter than 500 bp were excluded from the final assembly. Quality check of the assembly was performed using QUAST v5.2.0 (Gurevich et al. 2013) and checkM v1.2.2 (https://ecogenomics.github.io/CheckM/, Parks et al., 2015). The draft genome sequences of *Pseudomonas* sp. W2-17, *Pseudomonas* sp. W5-01, *Pseudomonas* sp. W5-36 and *Pseudomonas* sp. W2Oct36 have been deposited at DDBJ/EMBL/GenBank under accession numbers JAUFNS000000000.1, JAXCIP000000000.1, JAXCIQ000000000.1 and JAUERW000000000.1, respectively. Annotation of open reading frames (ORF) was performed with the NCBI Prokaryotic Automatic Annotation Pipeline (PGAAP).

### 2.9 Phylogenomic and receptor phylogeny analyses

A phylogenomic tree of 46 genomes belonging to the *P. lutea* clade was constructed using the Type (Strain) Genome Server TYGS (https://tygs.dsmz.de/). Phylogenetic analysis of formomarinobactin and marinobactin receptors from 134 selected strains was conducted using MEGA v12 (Kumar et al., 2024). Although over 500 receptor sequences were initially identified by BLAST, a selected set of representative sequences from reference strains was used to provide a tractable subset for analysis. The protein-based tree was inferred by Maximum Likelihood with the LG(+Freq) model (Le and Gascuel, 2008), selecting the highest log-likelihood topology (−31,242.43) from Neighbor-Joining and Maximum Parsimony starting trees; branch support reflects 1,000 bootstrap replicates (Felsenstein, 1985). Rate heterogeneity across 874 aligned amino acid sites was modeled with a 5-category discrete Gamma distribution (+G, parameter = 0.6174) and 17.39% invariant sites (+I). A complementary genome-based tree was constructed via Neighbor-Joining on a GGDC v3.0 formula 2 distance matrix (Meier-Kolthoff, 2022) derived from NCBI RefSeq assemblies (O’Leary et al., 2016).

### 2.10 Selection and siderophore production of *P. lutea* clade strains

From an updated *Pseudomonas* collection (∼ 1,000 strains) based on the one described in Matthijs et al., 2013 which is sequenced for the partial *rpoD* gene, strains belonging to the *P. lutea* clade were identified based on clustering with type strains of this clade. In addition to four *P. graminis* strains (P200/02, P265/09, P294/05 and P334/03) isolated concomitantly with the *P. graminis* type strain (Behrendt et al. 1999), four strains were obtained: *Pseudomonas* sp. W2-17, *Pseudomonas* sp. W5-01, *Pseudomonas* sp. W5-36 and *Pseudomonas* sp. W2Oct36. The latter four strains were isolated from a small Belgian river in year 2001 (W2Oct36, Matthijs et al., 2013) and 2006 (W2-17, W5-01 and W5-36, *unpublished*). Strains were characterized by CAS assay, and siderophores were purified and analyzed by LC-MS to determine the presence of formomarinobactins.

### 2.11 *In vitro* growth inhibition assay against Gram-positive and negative bacteria

The strains of the *P. lutea* clade were screened for their potential to produce iron-repressed molecules capable of inhibiting the growth of a panel of Gram-positive and Gram-negative bacterial strains (Table 1). Along with *P. graminis* LMG 21661^T^, eleven strains in the same *Pseudomonas* cluster were used to evaluate the conservation of the novel siderophore within the clade. Therefore, each *Pseudomonas* strain was grown overnight in 6 mL liquid CAA. The cultures were adjusted to 5 × 10^6^ cells, and 10 µL was inoculated in the center of 20 mL CAA plates and CAA plates supplemented with 50 µM FeCl_3_. The plates were then incubated at 28°C for 48h after which the cells were killed using chloroform vapors for 20 min. A soft agar overlay (7.5 g/L agar) mixed with a target strain adjusted at 5 × 10^6^ cells in 856 media was prepared. After evaporation of the chloroform, six mL of the overlay was uniformly poured on each plate and incubated overnight at 37°C. The assay was performed in triplicate for each condition.

The antimicrobial properties of the purified siderophores were evaluated against the same batch of test strains according to a slightly modified version of the previous assay. A soft agar overlay of each strain in 856 was prepared as described above and overlaid on 20 mL CAA and CAA + 50 µM iron plates. After solidification, a sterile cork borer was used to cut wells in the agar, which were then filled to the top of the well with 75 µl of siderophore fractions at 15 mg/mL in 70% MeOH. As control 70% MeOH was used. The plates were incubated overnight at 30 or 37^°^C, depending on the strain. The assay was performed in triplicate.

### 2.12 Formomarinobactin growth stimulation assays of non-producer and mutant strains

To identify strains with a putative formomarinobactin receptor, BLASTP searches were performed using the marinobactin receptor sequence of *P. graminis* as query against genomes of the genera *Pseudomonas*, *Ectopseudomonas*, *Stutzerimonas*, *Phytopseudomonas* and *Marinobacter*. Representative receptors with an E-value of 0 and a query coverage of ≥ 90 % were retained if they exhibited an amino acid identity of ≥ 60 % (or ≥ 48 % for *Marinobacter*). A representative of the clusteredNR results was chosen randomly for strains outside the *P. lutea* clade. Strains of interest, such as strains available in our in-house collection, were added to the selected candidates.

Siderophore-negative mutants derived from five of these identified strains, *P. simiae* WCS417, *P. capeferrum* WCS358, *Pseudomonas* sp. WCS374, *P. fluorescens* SBW25 and *P. rhodesiae* LMG 17764^T^ were used for growth stimulation assays, in addition to *P. bohemica* LMG 30182^T^, *P. kuykendallii* LMG 26364^T^ and *Phytopseudomonas argentinensis* LMG 22563^T^ (Table 1).

For the growth stimulation assays, pre-cultures of the strains grown overnight in CAA liquid medium were adjusted to 5 × 10^6^ cells/mL, then inoculated on 20 mL CAA plates supplemented with 2,2’-bipyridine. The optimal concentration of 2,2’-bipyridine was determined by testing a range from 100 to 600 µM, causing minimal background growth. A Whatman paper disk (Ø = 5 mm) was placed at the center of the plates, and 2 x 5 µL of purified siderophore was loaded onto it. Growth stimulation was assessed by recording growth halos of the mutant strains around the siderophore-loaded disks. Experiments were done twice.

### 2.13 Construction of a siderophore receptor mutant TWR49273 in *P. rhodesiae* LMG 17764^T^

To confirm that the siderophore receptors identified by BLAST analyses are formomarinobactin receptors, the putative receptor gene was deleted in the *P. rhodesiae* LMG 17764^T^ siderophore-negative mutant Prho^T^ΔpvdLΔtrbABC-44, hereafter referred to as Prho^T^-KO-44 (Brandt and Matthijs, *unpublished results*), which already carried clean deletions of biosynthetic siderophore genes, following a previously described method (Matthijs et al., 2016). In short, a markerless genomic deletion method by homologous recombination was used to achieve an in-frame deletion of the 11 kb internal fragment of the *pvdL* gene involved in biosynthesis of the chromophore of pyoverdine and of the almost complete *trbABC* operon of the siderophore triabactin (5 kb). Loss of pyoverdine and triabactin production were verified by LC-MS or CAS assay.

To delete the putative formomarinobactin TonB-dependent receptor, the 5′ and 3′ flanking regions were amplified and cloned in pUK21. After confirmation by sequencing the fragment was recloned in suicide pME3087, resulting in pME-TWR49273, and was integrated into the chromosome of Prho^T^-KO-44 by triparental mating using *E. coli* HB101/pME497 as the mobilizing strain, with selection for tetracycline and chloramphenicol resistant recombinants. Excision of the vector through a second crossing-over event took place following the enrichment of tetracycline-sensitive cells (Schnider-Keel et al. 2000). The mutant obtained was confirmed by PCR amplification and subsequent sequencing of the fragments (using verification primers, Supplementary Table S1). The list of the primers used to amplify the fragments is given in Supplementary Table S1, the constructed plasmids are given in Table 1.

### 2.14 Statistical analysis

One-Way ANOVA followed by Dunnett’s multiple comparison test for growth inhibition assays of the *P. lutea* clade strains and followed by posthoc Tukey correction for multiple comparisons were performed on growth inhibition assays of the purified formomarinobactins. A Brown-Forsythe and Welch ANOVA followed by Dunnett’s T3 test for multiple comparisons was done for RT-qPCR assays using Graphpad Prism version 8.0.1.

## 3 Results

### 3.1 *In silico* identification of an NRPS siderophore-mediated iron acquisition system in *P. graminis* LMG 21661^T^

Although *P. graminis* spp. do not produce pyoverdine (Behrendt et al. 1999; Butaitė et al. 2017; Peix et al. 2004), colonies of *P. graminis* LMG 21661^T^ exhibited a yellow halo on CAS medium, suggesting the synthesis of an alternative siderophore. Genome analysis identified a putative gene cluster for a NRPS siderophore which resided on a 37.5 kb fragment and included 11 open reading frames (ORFs) (Figure 1). Predictions showed that the necessary information for the regulation, biosynthesis and transport of a NRPS siderophore were present in the gene cluster.

**FIGURE 1.**
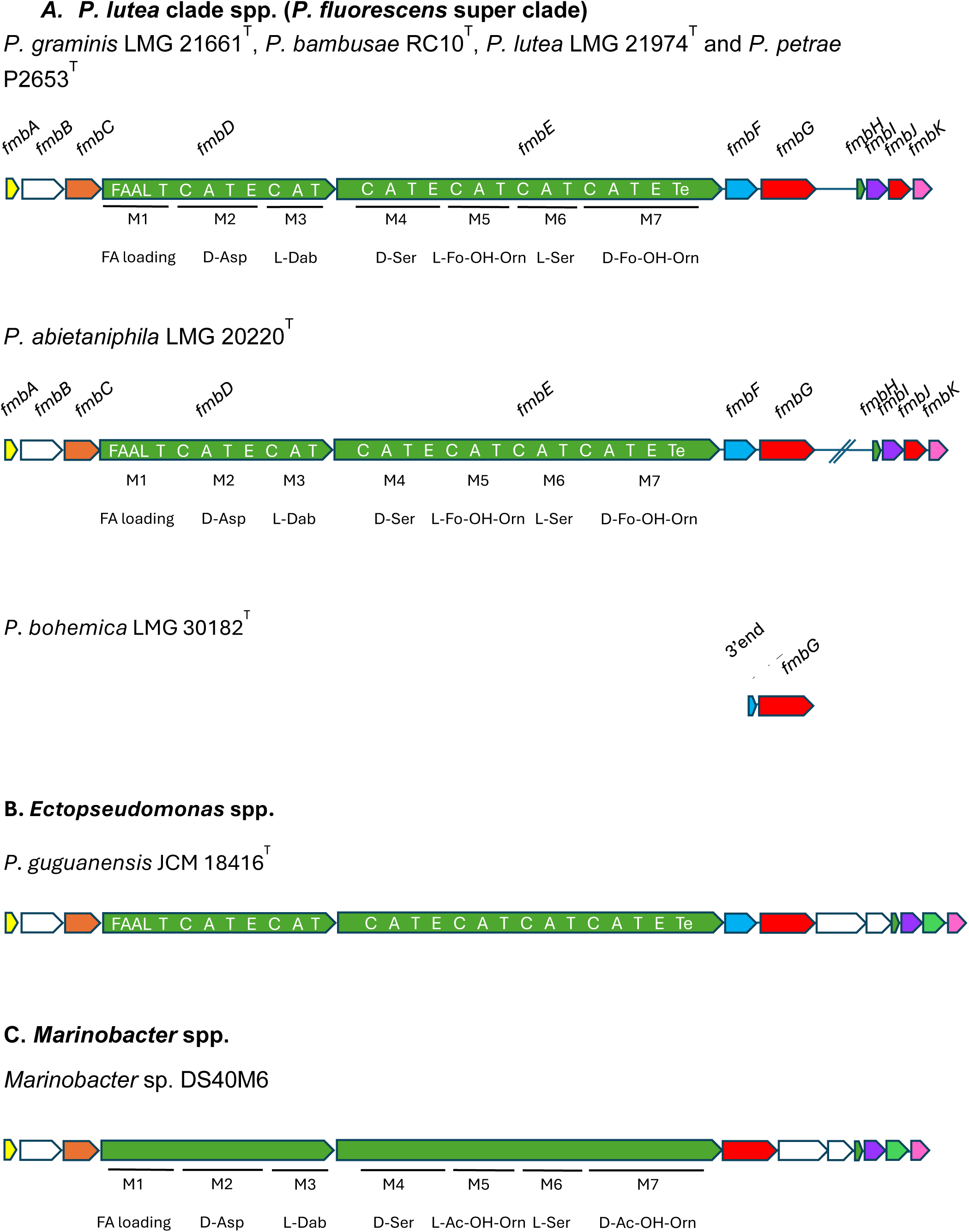
Schematic presentation of the gene cluster of formomarinobactin in strains of the *Pseudomonas* genus (**A**), the predicted marinobactin cluster of *Ectopseudomonas* spp. (**B**) and the marinobactin cluster of *Marinobacter* spp. (**C**). Gene names are indicated above the arrows. The organization of the modules and domains of the two NRPS, FmbD and FmbE, is shown on the arrows. Formo/marinobactin biosynthesis is predicted to begin with a N-terminal acyl AMP-ligase starter domain, which activates a fatty acid and initiates biosynthesis and to end with a thioesterase domain. C: condensation domain, A: adenylation domain, T: thiolation, E: epimerization domain. Predicted genes functions obtained by BLASTp and InterPro analyses: *fmbA*: RNA polymerase sigma-70 factor, ECF subfamily, *fmbB*: putative ATP-binding cassette transporter, *fmbC*: lysine N(6)-hydroxylase/L-ornithine N(5)-oxygenase family protein, *fmbD* and *fmbE*: non-ribosomal peptide synthases, *fmbF*: diaminobutyrate-2-oxoglutarate transaminase, *fmbG*: TonB-dependent receptor, *fmbH*: MbtH family NRPS accessory protein, *fmbI*: TauD/TfdA family dioxygenase, *fmbJ*: N(5)-hydroxyornithine transformylase PvdF and *fmbK*: siderophore-iron reductase FhuF. The homologue of FmbJ in *Ectopseudomonas* and *Marinobacter* spp. is a GNAT-family N-acetyltransferase.

The two NRPSs were predicted to assemble the peptide backbone. The organization of the modules and domains is shown in Figure 1. The length and composition of the peptide were inferred based on the domains of the NRPS. Six classical NRPS modules (C-A-T) were found, two in FmbD and four in FmbE, indicative of a 6-residues long peptide backbone. Adenylation domain specificity and the stereochemistry (D and L-forms, based on epimerization domains) were predicted (Supplementary S2), resulting in an <u>Asp</u> – Dab – <u>Ser</u> – (Fo-)OH-Orn – Ser – <u>(Fo-)OH-Orn</u> backbone (D-forms are underlined). In addition, a C-terminal thioesterase (TE) domain which catalyzes the release of the NRPS product was identified in the last protein FmbE. The gene cluster also encodes a MbtH-like protein (FmbH) which is thought to assist many NRPSs by promoting proper folding, stability, and activity of the synthetases (Felnagle et al. 2010).

The first domain of the predicted FmbD protein presented homology with members of the fatty acyl-AMP ligase family (FAAL, cd05931, e = 0) (Figure 1). This domain may activate and introduce long chain fatty acids in both polyketide and NRPS peptide biosynthesis by activation and loading of fatty acids to ACP domains (Patil et al. 2021). The presence of this domain led to the hypothesis that the siderophore is amphiphilic.

Two non-proteinogenic amino acids, 2,4-diaminobutanoic acid (Dab) and (formylated) N5-hydroxyornithine [(Fo-)HO-Orn], were proposed as components of the peptide of the siderophore. L-Dab is probably synthesized from aspartate *ß*-semialdehyde by diaminobutyrate-2-oxoglutarate transaminase (FmbF). L-ornithine N5-oxygenase (FmbC) converts L-ornithine to N_5_-hydroxyornithine. Although the software had difficulty differentiating between non-formylated and formylated hydroxyornithine, the presence of an N(5)-hydroxyornithine transformylase PvdF (FmbJ) in the cluster suggests the hydroxyornithine is likely in the formylated state (McMorran et al. 2001; Ravel and Cornelis 2003). These modifications generate hydroxamate groups that chelate Fe(III) via oxygen atoms, creating stable iron complexes. Formylated *N^5^*-hydroxyornithines are characteristic for pyoverdines and contribute strongly to iron chelation (Meyer 2000). An additional coordination site is probably a hydroxy-carboxylic group formed by the hydroxylation of aspartate, catalyzed by the aspartyl-β-hydroxylating tailoring enzyme FmbI. The last gene, *fmbK*, coded for a protein that is homologous to FhuF, the putative iron-ferrioxamine B reductase in *E. coli*, which is proposed to reductively release ferrous iron from ferric-siderophore complexes. FmbK is thus likely involved in the release of iron from internalized ferric-siderophore that then becomes available for cellular use. Finally, the uptake of the ferric-siderophore complex probably occurs through the TonB-dependent receptor FmbG.

Overall, the gene cluster indicates production of a marinobactin-like siderophore, with an N(5)-hydroxyornithine transformylase replacing a GNAT-family N-acetyltransferase (Figure 1), predicting a formylated rather than acetylated siderophore.

### 3.2 Siderophore gene cluster transcription is regulated by iron availability

To evaluate if the *fmb* genes from *P. graminis* LMG 21661^T^ are functional and regulated by iron, expression of the first NRPS gene *fmbD* and the TonB-dependent receptor *fmbG* was verified by means of RT-qPCR under iron limited and iron-rich conditions. A preliminary assay of six bacterial cultures supplemented with increasing iron concentrations (0, 0.5, 1, 2, 4 and 8 µM FeCl_3_) showed that the *fmbD* and *fmbG* genes were expressed in the absence of iron and repressed by low amounts of Fe^3+^ in the medium. Indeed, exposure to 0.5 µM resulted in complete repression of *fmbD* expression and an approximately one-third reduction in *fmbG* expression (data not shown). Iron concentrations were refined (0, 0.05, 0.1, 0.3, 0.5, 0.7 µM FeCl_3_) to assess more precisely the iron-dependent regulation of gene expression (Figure 2). A concentration of 0.3 μM FeCl_3_ significantly reduced *fmbD* transcription (*p-value* <0.05), the first NRPS gene while the receptor transcription decreased by approximately one third, with complete suppression observed at 0.7 µM FeCl_3_ (*p-value* <0.01). These results indicate that the cluster is iron regulated. The effect of zinc and nickel on the expression of *fmbD* and *fmbG* was also tested, no effect was observed (Supplementary Figure S1) excluding a role as zincophore or nickelophore for formomarinobactin.

**FIGURE 2.**
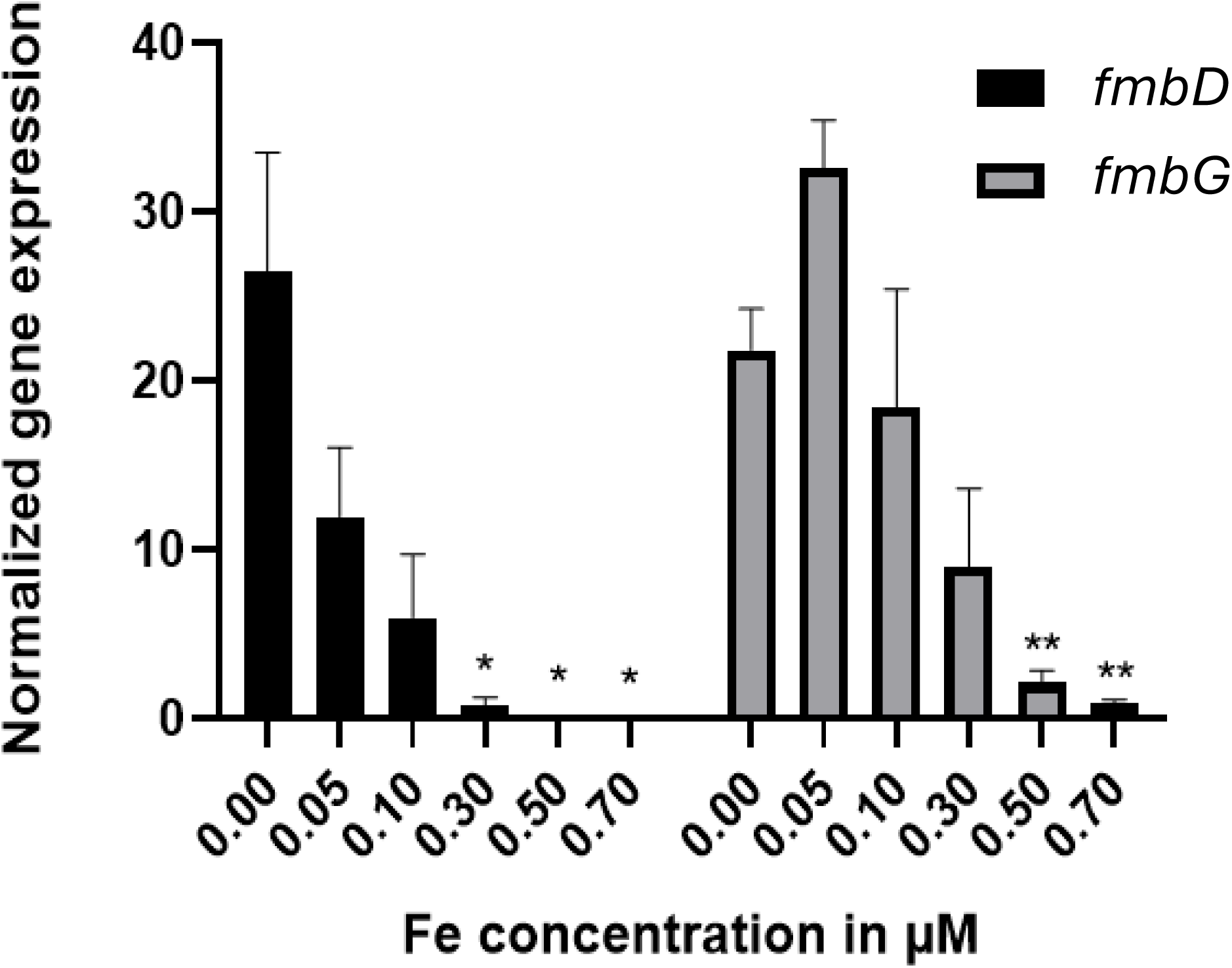
Relative expression of *fmbD* and *fmbG* genes with increasing Fe concentrations normalized against reference gene *nadB*. Experiments were repeated six times. Values correspond to mean ± SEM. A Brown-Forsythe and Welch ANOVA analysis followed by Dunnett’s T3 test for multiple comparisons was done. * p <0.05, ** p <0.01.

### 3.3 Structure of marinobactin-like siderophore suite produced by *P. graminis* LMG 21661^T^

Six peaks were successfully purified as separate molecules by preparative HPLC, and their purity was assessed by LC-MS.

A LC-HRMS full scan analysis was conducted to obtain the precise mass of each molecule. These were determined as *m/z* 874.4177 (F_A_), 876.4379 (F_B_), 902.4542 (F_C_), 904.4655 (F_D_), 930.4878 (F_E_) and 932.5002 (F_F_) for the [M+H]^+^ ions (Figure 3). According to their *m/z* values and close retention times, they were hypothesized to be a suite of siderophores. Furthermore, mass differences of 2 Da between *m/z* 874 and 876, 902 and 904 and, 930 and 932, seemed to suggest 2H corresponding to an unsaturation while the mass differential between *m/z* 876 and 902 as well as 904 and 930 of 26 Da, are probably the result of C_2_H_2_, so methylene groups. The mass spectrum consisted of the [M+H]^+^ monoisotopic ion as well as the doubly charged [M+2H]^2+^ ion (Supplementary Figure S2). Finally, the molecular formulae of the siderophores were also determined by considering isotopic distribution.

**FIGURE 3.**
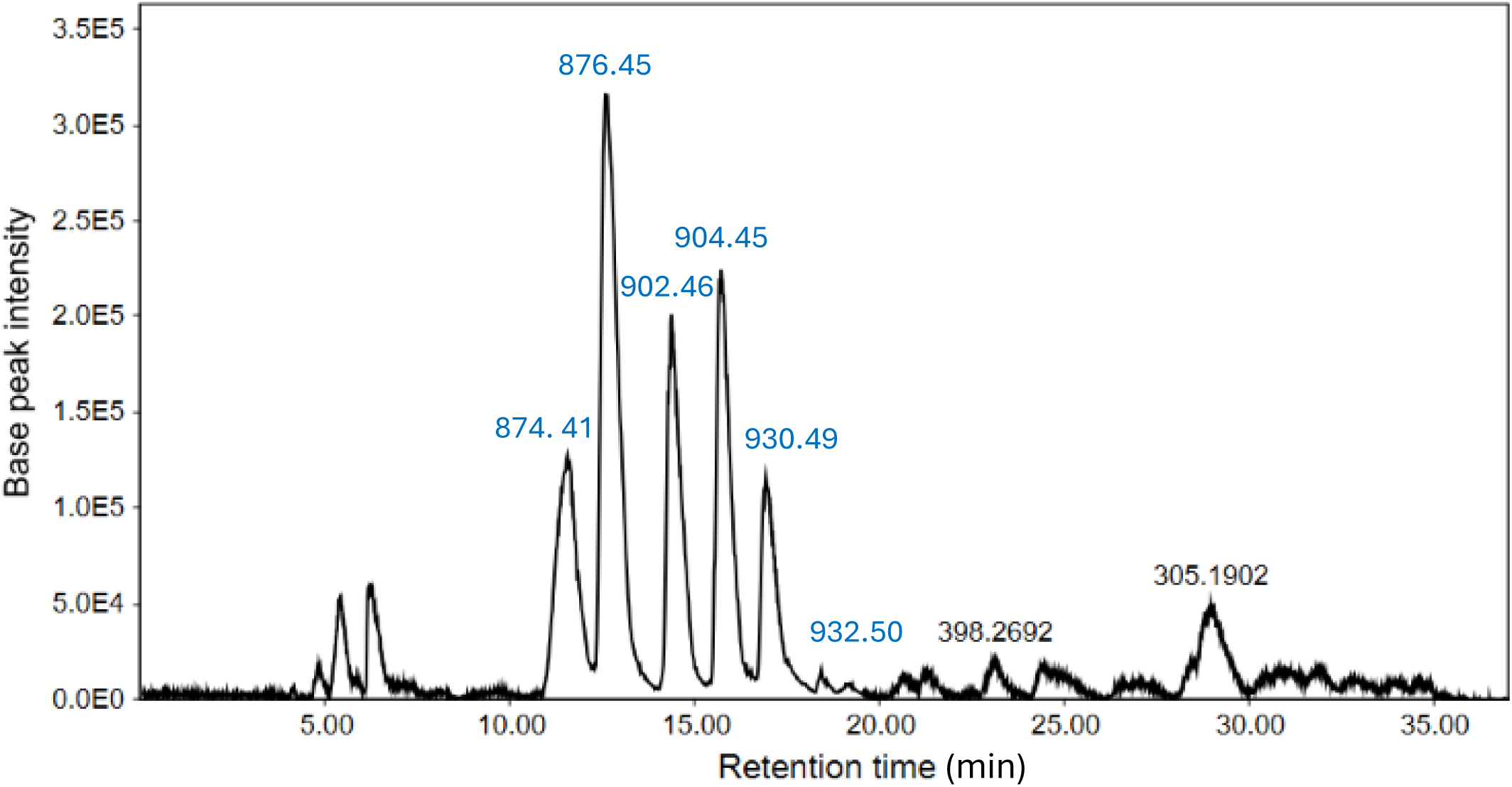
LC-HRMS chromatogram of semi-purified formomarinobactin-rich fraction in positive mode. Six peaks represent *m/z* values of each formomarinobactin variant.

The LC-MS/MS analysis showed similar fragmentation patterns (neutral losses) for all siderophores at both 35 eV (Supplementary Figure S3) and 55-65 eV indicating similar structures. Moreover, fragmentation at 55-65 eV, involved the loss of a water molecule (18 Da), then CO_2_ (44 Da) characteristics of loss of the carboxylic group on the C-terminal. This is followed by the loss of a fragment between 152 Da (for *m/z* 874 [M+H]^+^) and 210 Da (for *m/z* 932 [M+H]^+^) from the N-terminal resulting in a common *m*/*z* 660 Da fragment. Also, the same fragmentation peaks were observed below *m/z* 264/265 denoting a common core (Supplementary Figure S4) and enabling the elucidation of the amino acids present in the peptide sequence. Many of these peaks are shared with marinobactins, namely *m/z* 86, 150, 170, 211, 219, 237, 255 et 265 (Reitz et al. 2025; Zhang 1999). In fact, main differences with the fragmentation of marinobactins concerned modifications of the ornithine residue, instead of a loss of 190 and 172 Da for N-acetyl N-hydroxyornithine and N-acetyl N-hydroxyornithyl, respectively, losses of 176 and 158 Da were observed corresponding to N-formyl N-hydroxyornithine and an N-formyl N-hydroxyornithyl residue, respectively (Figure 4). The formylated ornithine residues were alternated with serine (seryl groups), observed by a loss of 87 Da, similarly to marinobactins, followed by the loss of diaminobutyric acid (100/101 Da). As described by Zhang 1999, fragments corresponding to the loss of Dab and Asp/Asp(OH) as well as the fatty acid tail are less evident, especially in the formomarinobactin variants of higher masses. Nonetheless, fragmentation of *m/z* 904 [M+H]^+^ at 55eV was strongly similar to that recorded for marinobactin A according to Reitz et al. 2025, suggesting that the molecules have highly similar structures. Marinobactin A has a *m/z* of 932 [M+H]^+^ in positive mode, the modification of its Ac(OH)Orn groups to Fo(OH)Orn, would lead to a *m/z* of 904. The results are consistent with the *in silico* analysis, predicting a marinobactin-like siderophore composed of a common peptidic backbone and a fatty acid chain of varying length (Figure 4). The presence of a hydroxamate-type siderophore was further confirmed by TLC analysis (Supplementary Figure S5).

**FIGURE 4.**
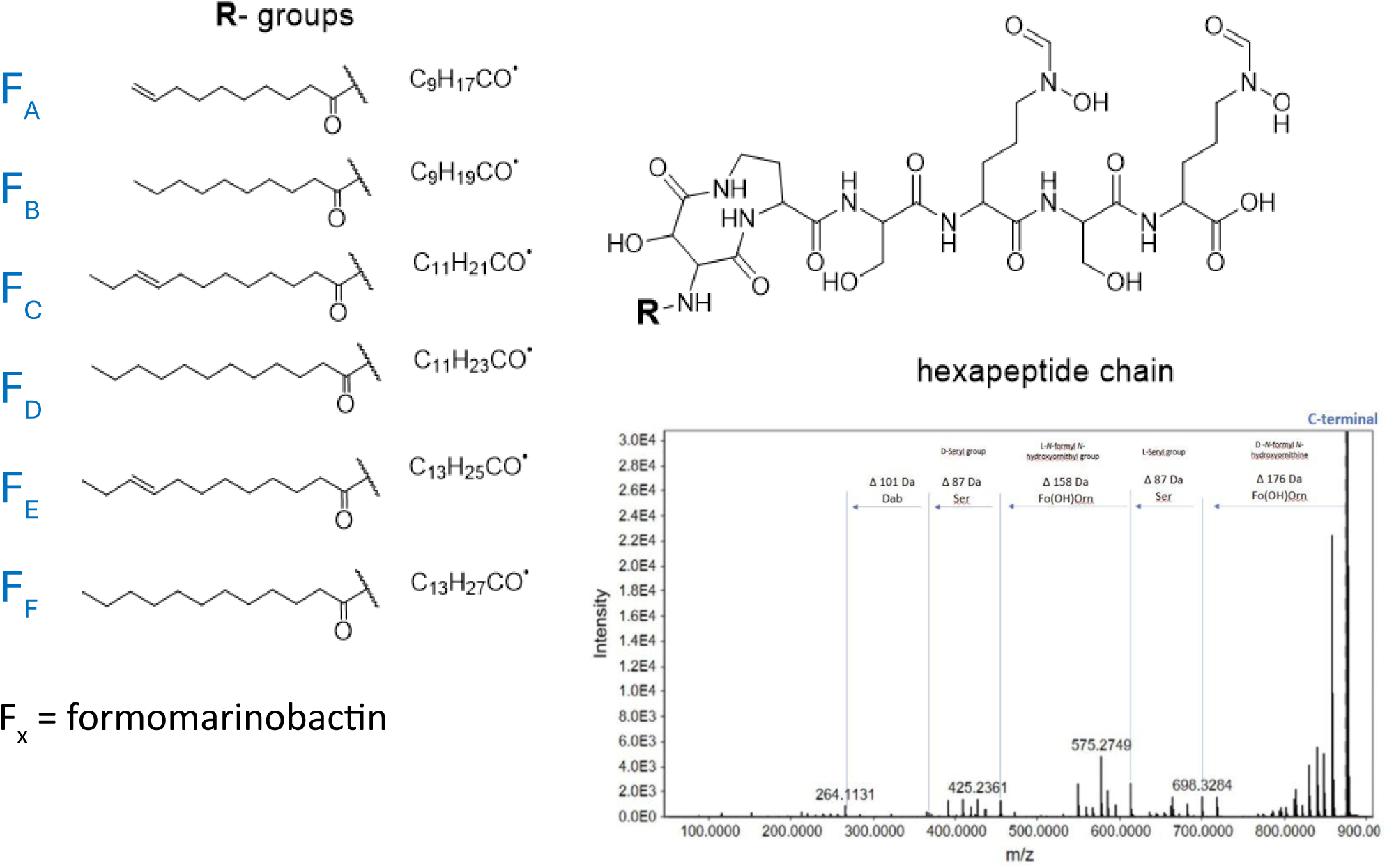
Formomarinobactin structure and fragmentation of peptide chain from COOH-terminal. Example with 874 *m/z*.

In light of the sole modification on the ornithine residue, we propose naming our series of siderophores formomarinobactins: *m/z* 874 Formomarinobactin A, *m/z* 876 Formomarinobactin B, *m/z* 902 Formomarinobactin C, *m/z* 904 Formomarinobactin D, *m/z* 930 Formomarinobactin E and *m/z* 932 Formomarinobactin F. Formomarinobactin D (F_D_) thus has a lauryl group (C12:0) for fatty acid tail, as also observed from the common fragment at 660 Da. F_F_ has a tetradecanoic acyl group (C14:0), F_E_ likely has a myristoleic acyl group (C14:1) and F_B_ a decanoic acyl group (C10:0). F_A_ and F_C_ both likely carry an unsaturation at Δ9 as (C10:1) and (C12:1), respectively.

### 3.4 Formomarinobactins are the characteristic siderophores of the *P. lutea* clade of the *Pseudomomas* genus

The studied *P. graminis* strain belongs to the *P. lutea* clade (*P. fluorescens* superclade), a taxonomic group that currently includes six described species: *P. abietaniphila*, *P. bambusae* (not validly publ.), *P. bohemica*, *P. graminis*, *P. lutea* and *P. petrae* (Figure 5). Phylogenomic analysis resolved the *P. lutea* clade into two well-supported subclades, A and B, containing 13 and 11 (putative) species, respectively, suggesting 18 predicted novel species in addition to the six described. dDDH analysis indicated that *P. bambusae* may represent two subspecies (Figure 5); its dDDH value with *Pseudomonas* sp. SLFW was 72.3%, near the lower boundary of the commonly accepted species threshold.

**FIGURE 5.**
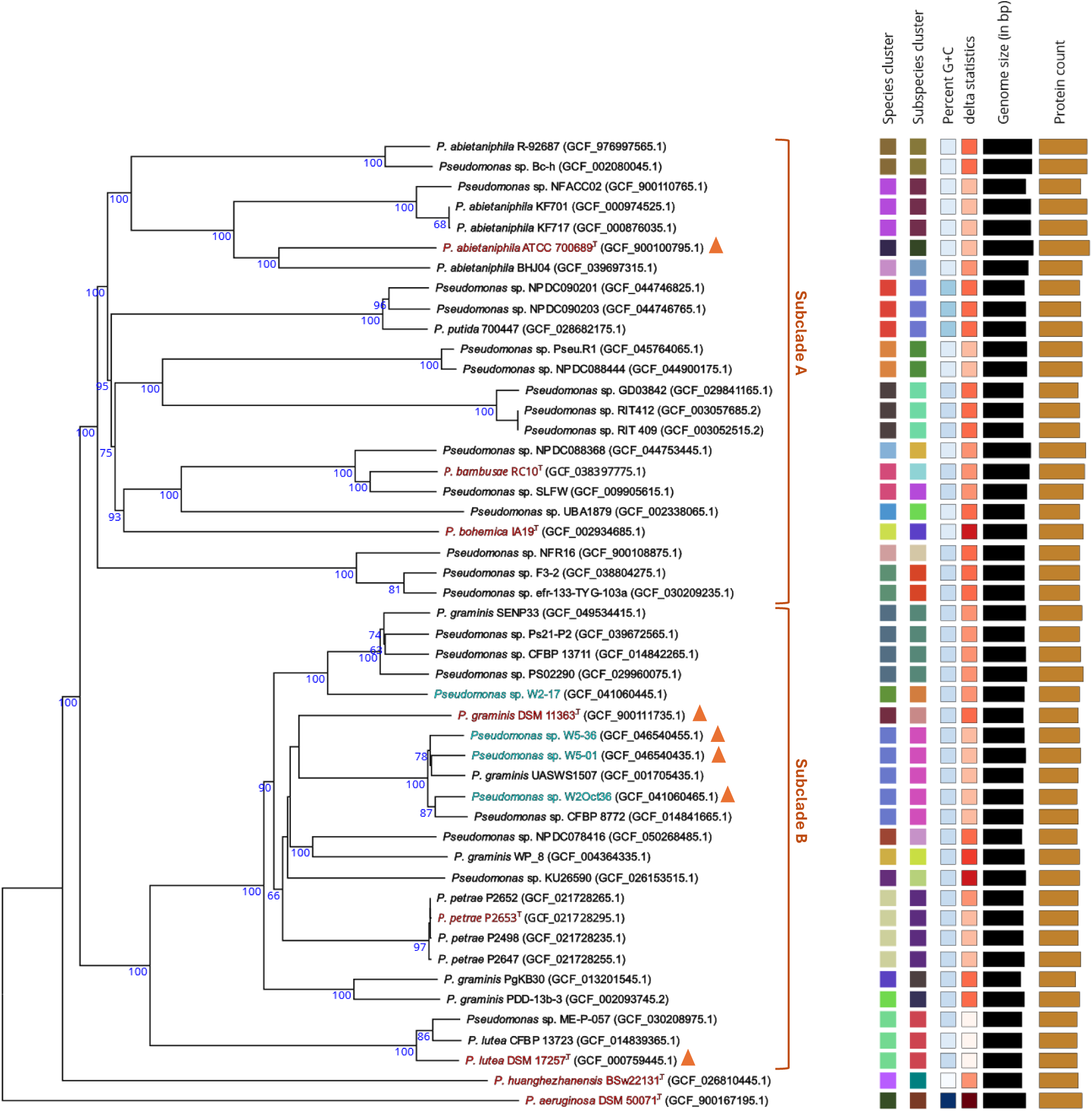
Phylogram generated using the TYGS platform (https://tygs.dsmz.de) (Meier-Kolthoff and Göker, 2019). Type strains are shown in red, genome-sequenced strains analyzed in this study are shown in blue, and orange triangles indicate experimentally confirmed formomarinobactin production.

Although *P. huanghezhanensis* shows its highest genomic similarity to members of the *P. lutea* clade (Ge et al., 2024), the dDDH values are very low (< 24.1%), and it forms a long-branched, genomically distant lineage sister to the *P. lutea* clade (Figure 5). Due to this uncertain taxonomic position, *P. huanghezhanensis* was excluded from downstream analyses.

Genome mining revealed that all strains of the *P. lutea* clade, apart from the type strain of *P. bohemica* (Figure 1), harbor a formomarinobactin biosynthetic gene cluster. In some genomes, this cluster is split, with the *fmbH-K* genes located distantly from the core biosynthesis genes, as observed, for example, in the *P. abietaniphila* type strain (Figure 1).

LC-MS analysis of the available type strains of the *P. lutea* clade confirmed that *P. graminis*, *P. lutea* and *P. abietaniphila* exhibited six peaks with *m/z* 874.7, 876.7, 902.7, 904.7, 930.7 and 932.7. In addition, formomarinobactin production was detected in four additional *P. graminis* strains whose genomes have not been sequenced (*P. graminis* P200/02, P294/05, P265/09 and P334/03), as well as in four strains isolated from a small river (*Pseudomonas* sp. W2Oct36, W2-17, W5-01, and W5-36) (Table 1). These results, combined with genome mining, indicate that formomarinobactin production is conserved across the *P. lutea* clade.

Outside the *P. lutea* clade, a formomarinobactin gene cluster was identified in *P. huanghezhanensis* BSw22131, a pyoverdine-negative strain. The cluster is flanked by an IS3-family transposase in the intergenic region between the receptor and accessory genes, with a site-specific integrase, phage tail tape measure protein, and IS4-family transposase located upstream. These genomic features indicate recent mobility and suggest horizontal acquisition and genomic instability, which may facilitate biosynthetic gene loss or the generation of variants.

Putative marinobactin gene clusters, identified by the presence of a GNAT-family N-acetyltransferase and absence of N(5)-hydroxyornithine transformylase, are rare within the genus *Pseudomonas*. Only a single cluster was identified, in *P. oryzihabitans* USDA-ARS-USMAR, a pyoverdine non producer. In contrast, marinobactin clusters were detected outside the genus, including in *Phytopseudomonas argentinensis* SA190 and multiple *Ectopseudomonas* strains (Figure 1). Similar to the formomarinobactin cluster, and unlike *Marinobacter*, these gene clusters are self-sufficient for 2,4-diaminobutyrate (Dab) biosynthesis.

The marinobactin cluster has previously been investigated in *Ectopseudomonas mendocina* ymp (formerly *Pseudomonas mendocina*), where it was shown to contribute to iron acquisition; however, the chemical structure of the corresponding siderophore has not yet been elucidated (Awaya and DuBois 2008).

### 3.5 Growth stimulation assays of putative formomarinobactin TonB-dependent receptors

BLAST analysis revealed that homologs of formomarinobactin/marinobactin TonB-dependent receptors, in the absence of the corresponding biosynthetic gene cluster, are widely distributed across the *P. fluorescens* superclade. Over 1000 receptor homologs were identified over multiple genera. Because a full phylogenetic reconstruction was beyond the scope of this study, a representative set of 134 genomes was analyzed (Figure 6). Gene- and genome-based phylogenies were constructed to compare receptor relationships with overall genomic relatedness and tree congruence was qualitatively assessed.

**FIGURE 6.**
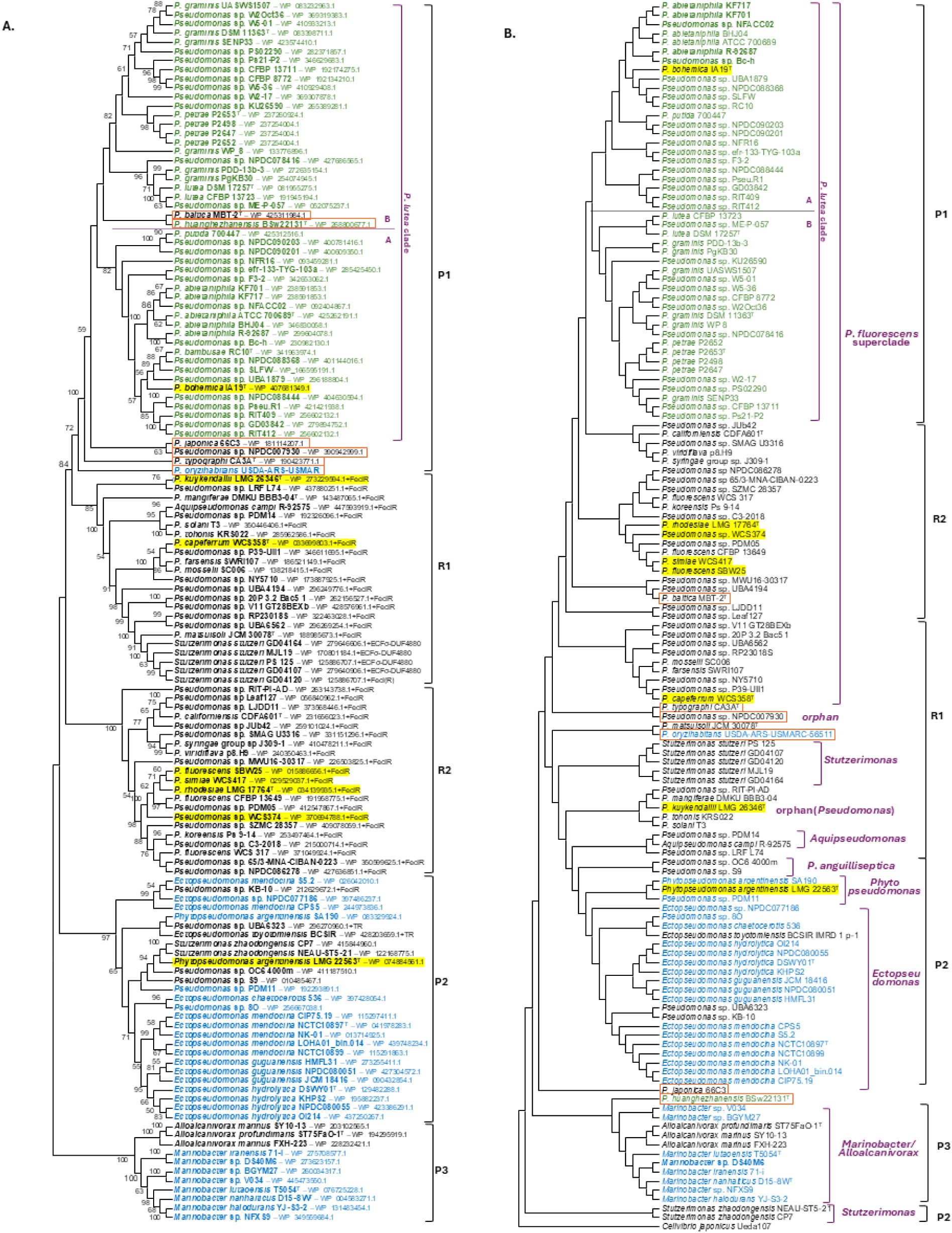
Comparative phylogeny (topology-only) of formomarinobactin and marinobactin receptors and host genomes. **A**: Maximum Likelihood tree of 134 representative receptors (subset of >1,000 BLAST hits), inferred from 874 aligned amino acid sites using the LG(+F+G+I) model in MEGA v12 (1,000 bootstrap replicates; values >50% shown). **B**: Neighbor-Joining tree of the same 134 strains based on GGDC v3.0 formula 2 intergenomic distances from NCBI RefSeq assemblies (*Cellvibrio japonicus* outgroup); no branch supports shown due to distance-matrix method. Color coding: formomarinobactin producers (green), marinobactin producers (blue), receptor-only strains (black). Yellow highlight: growth stimulation assay strains. Orange squares: topologically incongruent strains. Cell-surface signaling upstream of receptors in non-producers: FecIR (Fec system σ⁷⁰+FecR), ECFσ–DUF4880 (ECFσ⁷⁰+DUF4880). Parentheses: partial genes; TR: siderophore transport genes.

The Maximum Likelihood receptor phylogeny resolved three producer-associated subclades (P1-P3) and two receptor-only subclades (R1-R2), each supported by bootstrap values >70% (Figure 6A). Subclade P1 comprised formomarinobactin producing *Pseudomonas* strains, whereas P2 and P3 contained marinobactin producers from *Ectopseudomonas* and *Marinobacter*, respectively. Within producer-associated subclades, several smaller receptor-only lineages were present. Clade R2 was composed of *Pseudomonas* spp., while R1 included *Stutzerimonas*, *Aquipseudomonas* and orphan *Pseudomonas* strains. Receptors in R1 and R2 were preceded by cell-surface signaling (CSS) genes (ferIR loci) or, in *Stutzerimonas*, by an ECFσ–DUF4880 module, whereas incongruent, or marinobactin-producing strains, lacked identifiable CSS systems.

The complementary GGDC-based Neighbor-Joining genomic tree revealed several taxonomic inconsistencies relative to the receptor phylogeny. For example, within the formomarinobactin producing P1 clade, nested the receptor-only *P. baltica* type strain (*P. rhizosphaerae* clade) and the predicted formomarinobactin producing *P. huanghezhanensis* BSw22131^T^ (taxonomic position not clear) (Figure 6A). Adjacent to the *P. lutea* clade were receptors from *P. japonica* 66C3, *Pseudomonas* sp. NPDC007930, *P. typographi* CA3A^T^, and *P. oryzihabitans* USDA-ARS-USMAR, which were distantly related to the *P. lutea* clade in the genome-based phylogeny (Figure 6B).

In the *P. lutea* clade the core formomarinobactin biosynthetic gene cluster was located immediately upstream of the receptor gene, except in *P. bohemica*, which harbored only the receptor preceded by a truncated 3’ end of *fmbE*. Growth stimulation assays showed that *P. bohemica* grew in the presence of purified formomarinobactin (Figure 7A), confirming receptor functionality and substrate specificity.

**FIGURE 7 A-D.**
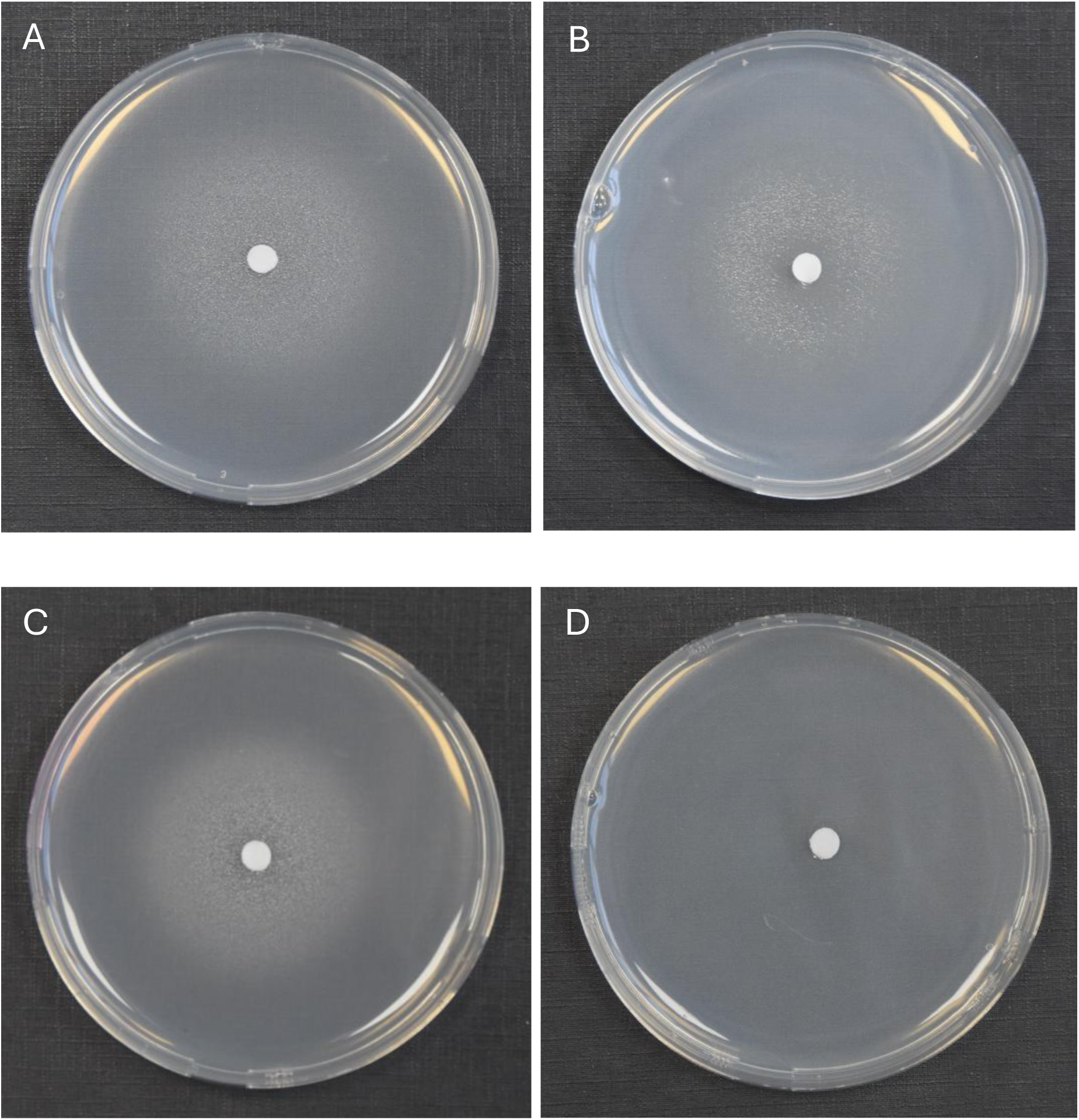
Formomarinobactin growth stimulation assays on CAA medium supplemented with the iron chelator 2,2′-bipyridine. (**A**) *P. bohemica* LMG 30182^T^ (500 µM), (**B**) *Phytopseudomonas argentinensis* LMG 22563^T^ (formerly classified as *Pseudomonas argentinensis*) (200 µM), (**C**) Siderophore-negative mutant Prho^T^KO-159 of *P. rhodesiae* LMG 17764^T^ (400 µM) and (**D**) Prho^T^KOΔTWR49273-159 (lacking the formomarinobactin receptor) (400 µM).

To test whether receptor-only strains outside the *P. lutea* clade could utilize formomarinobactin as an iron source, growth stimulation assays were performed with selected receptor-only strains (highlighted in yellow in Figure 6) from the subclades P2, R1 and R2. Siderophore-deficient mutants of *P. fluorescens* SBW25, *P. simiae* WCS417, *P. rhodesiae* LMG 17764^T^, *Pseudomonas* sp. WCS374 (all *P. fluorescens* subclade) and *P. capeferrum* WCS358 (*P. putida* clade), in addition to *P. kuykendallii* LMG 26364^T^ (orphan clade) all grew around formomarinobactin-loaded disks (Supplementary Figure S7A-D), confirming active, functional receptors.

To directly demonstrate that the predicted TonB-dependent receptors mediate formomarinobactin uptake, the receptor gene was deleted in the siderophore-deficient mutant Prho^T^-KO-44, yielding Prho^T^-KOΔTWR49273-159. The double mutant failed to grow in the presence of formomarinobactin (Figure 7C-D), demonstrating that the receptor is essential for formomarinobactin-dependent iron uptake.

### 3.6 Iron-mediated growth inhibition of *P. lutea* clade strains against target microorganisms

The strains of the *P. lutea* clade showed clear antagonistic activity against two Gram positive bacteria, *S. capitis* I and *B. zanthoxylo* RSN02 (Figure 8 and 9, and Supplementary Figure S7). The antibacterial activity observed was iron dependent as its supplementation at 50 µM in the medium resulted in the loss of inhibition zones. Two Gram negative strains, *P. aeruginosa* LMG 1242^T^ and *Enterobacteriaceae* sp. DV4951, were partially inhibited by the strains, visible as a reduced lawn density (Figure 8 and 9, and Supplementary Figure S7). This indicates that both target strains are stressed but can still grow in the presence of the exometabolites under the tested conditions. No inhibition was once again observed in iron-rich conditions. Conversely, *P. bohemica* LMG 30182^T^, which does not produce formomarinobactin, was the sole exception in these growth inhibition assays; it showed no inhibition against any of the Gram-positive and Gram-negative bacterial strains, whether in iron-rich or iron-limited media (Figure 8 and 9). The iron-dependent repression of the growth inhibition, together with its absence in *P. bohemica* supports the involvement of formomarinobactins in the observed growth inhibition.

**FIGURE 8.**
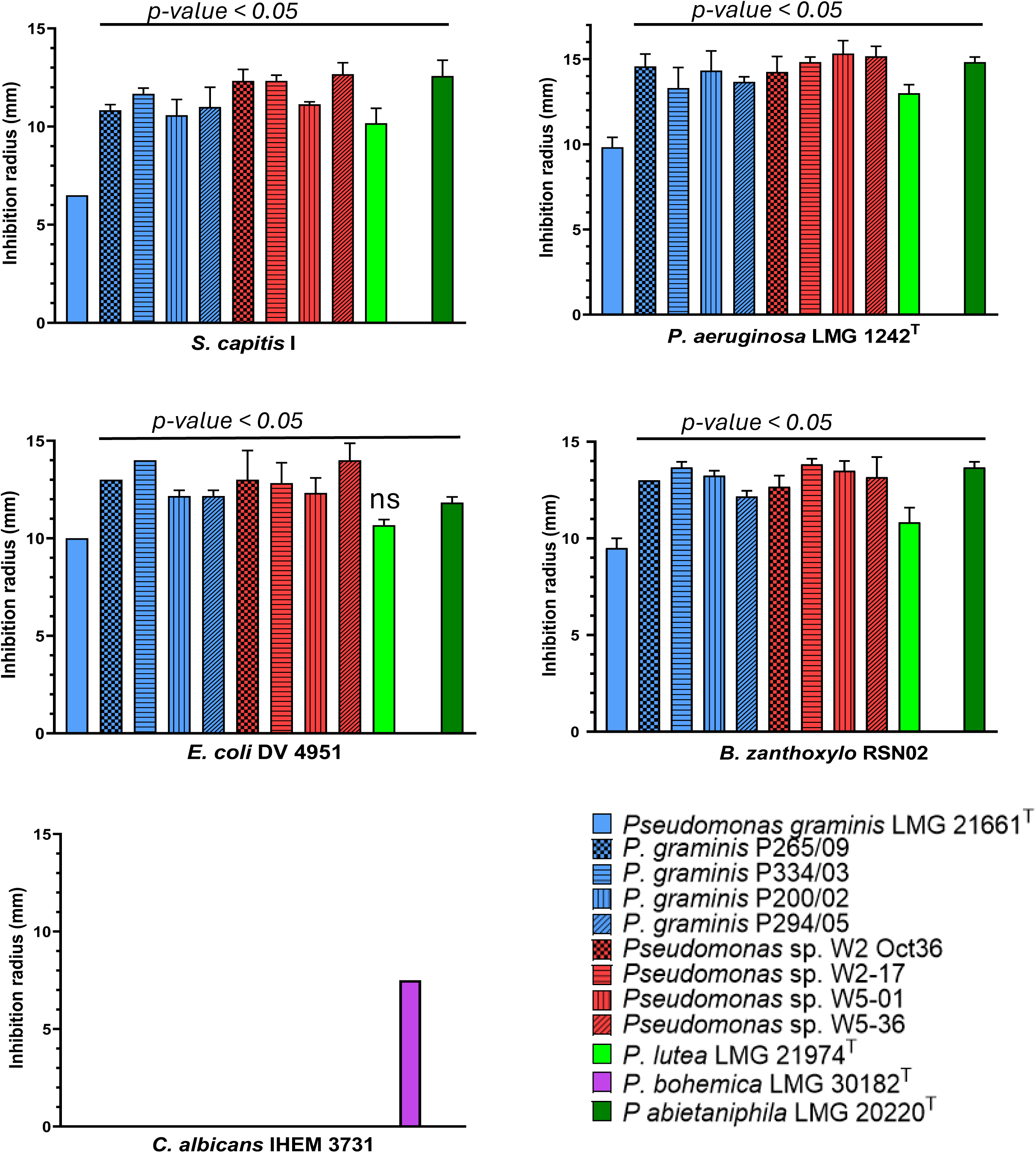
Antagonism of *P. lutea* clade strains against clinical and environmental strains. Experiments were done in triplicate. Values correspond to mean ± SD. One-Way ANOVA followed by Dunnett’s multiple comparisons test. Significance level at *p-value* < 0.05 compared to LMG 21661^T^ for all *P. lutea* G strains against antagonistic bacterial strains except LMG 21974^T^ against *E. coli (ns = non significant)*.

**FIGURE 9.**
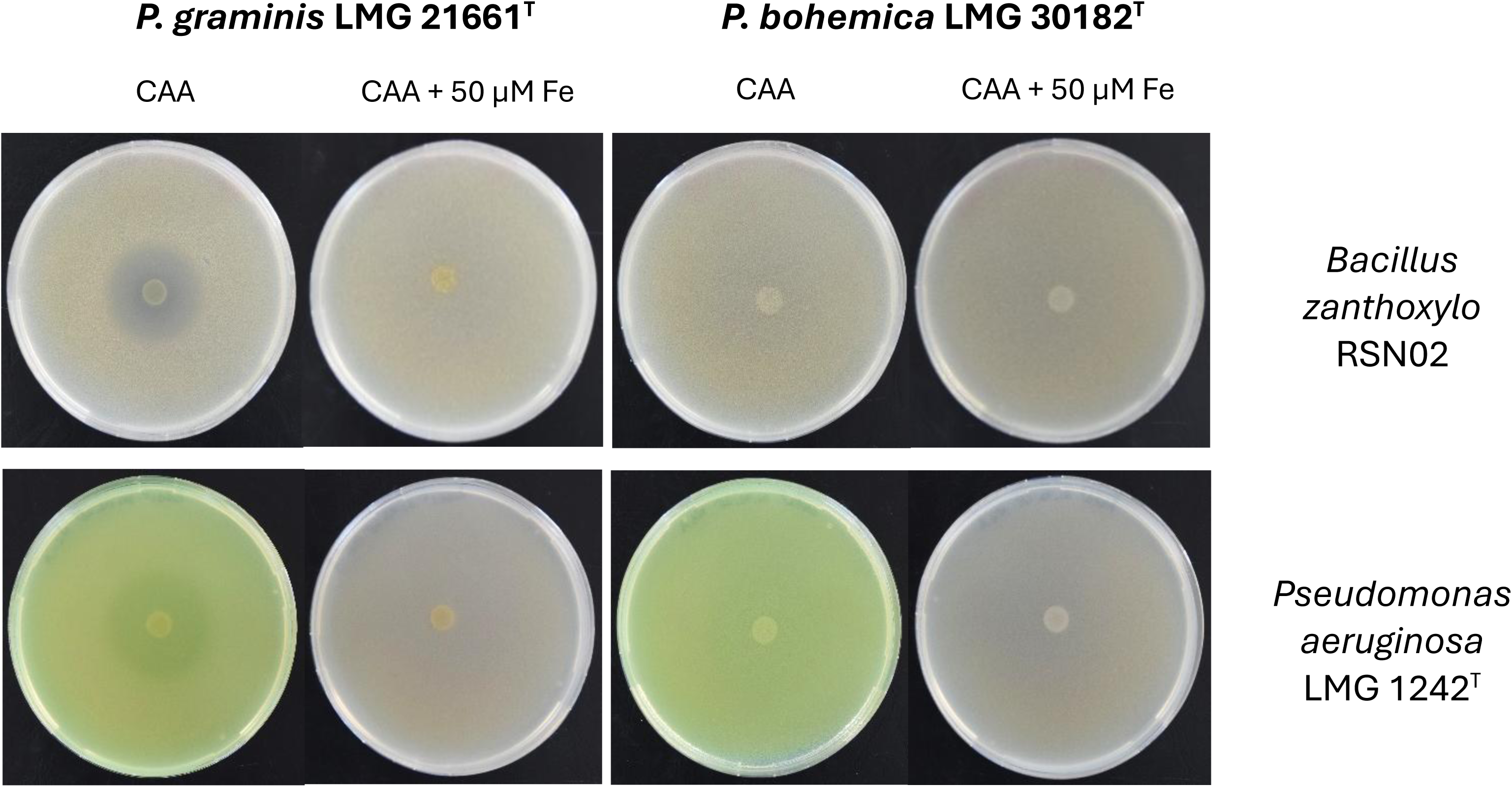
Growth inhibition of *P. graminis* LMG 21661^T^ (left column) and *P. bohemica* LMG 30182^T^ (right column) against the Gram-positive *B. zanthoxylo* RSN02 and the Gram-negative *P. aeruginosa* LMG 1242^T^. Images are linearly adjusted for contrast (+20%) for presentation. The results of the growth inhibition against the other strains can be found in **Supplementary Figure S7**.

Finally, *E. hirae* was the only clinical strain against which none of the *P. lutea* clade strains were active. Interestingly, none of the strains showed anti-yeast activity against *Candida albicans*, except for *P. bohemica* LMG 30182^T^ (7.5 mm).

Furthermore, in each assay, the inhibition zones of the wild-type, *P. graminis* LMG 21661^T^, were significantly (*p-value* < 0.05) smaller than those of the other producer strains, highlighting either a lower yield of siderophore or the secretion of other iron-suppressed antibacterial substances by the *P. lutea* clade strains (Figure 8).

### 3.7 Growth inhibiting potential of *P. lutea* clade strains is mediated by amphiphilic siderophores, formomarinobactins

The results obtained for the isolated molecules mirrored those of the strains with a clear growth inhibition against *B. zanthoxylo* RSN02 and *S. capitis* I (Figure 10). All 6 molecules showed a similar range of inhibition. The inhibition zones were greater in iron-poor than iron-rich media, however they were distinctly observed in both conditions. These findings further imply that the growth inhibition of the strains was mediated by iron-dependent molecules. The only exception was *Enterobacteriaceae sp.* DV 4951 which was not inhibited by the purified molecules, perhaps suggesting the presence of other iron-mediated anti-microbial substances or that their activity against DV 4951 was only significant in combination rather than as individual molecules.

**FIGURE 10.**
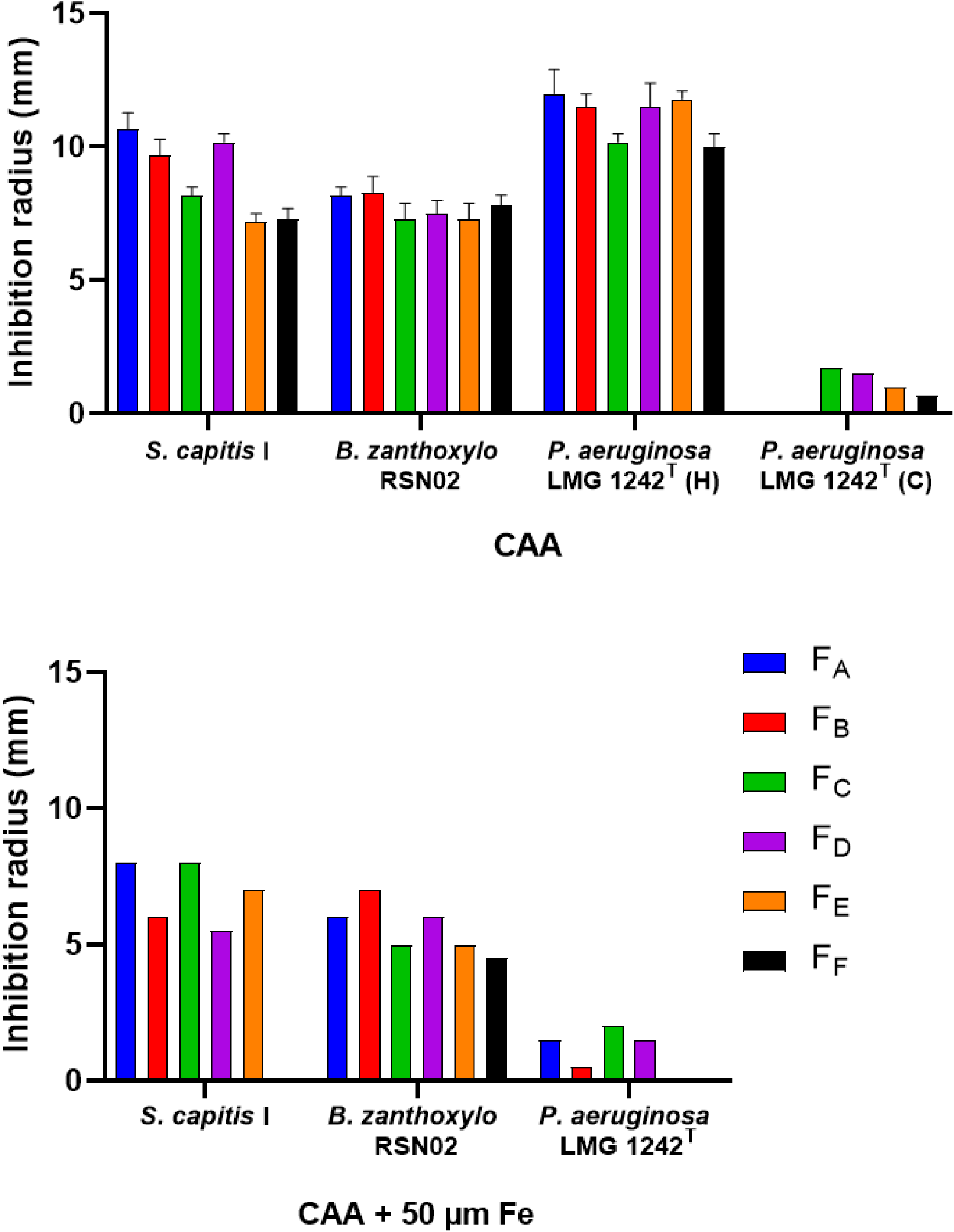
Growth inhibition of purified formomarinobactins F_A_ (*m/z* 874), F_B_ (*m/z* 876), F_C_ (*m/z* 902), F_D_ (*m/z* 904), F_E_ (*m/z* 930) and F_F_ (*m/z* 932) against clinical and environmental strains in iron-limited (left) and iron-supplemented (right) CAA medium. H = halo, for reduced lawn density; C = clear antagonism, for radius devoid of growth. Experiments were done in triplicate. Values correspond to mean ± SD. One-Way ANOVA followed by posthoc Tukey correction for multiple comparisons was applied and significance levels were defined as *p<0.05, **p<0.01, ***p<0.001, ****p<0.0001.

*C. albicans* and *E. hirae* were not inhibited by the molecules tested, confirming that the lack of growth inhibition observed in the presence of the *P. lutea* clade strains was due to the lack of activity of the siderophores against these two strains.

*P. aeruginosa* showed a unique inhibition pattern. Indeed, consistent with results from the growth inhibition of the strains, a reduced lawn density halo was observed in the presence of the different siderophores suggesting partial growth inhibition. Additionally, clear growth inhibition was observed for F_C_ – F_E_ (Supplementary Figure S8) with a small inhibition zone devoid of growth. This inhibition zone was also observed in the presence of iron, without the presence of partial growth. Such results highlight that the growth inhibition of formomarinobactins against *P. aeruginosa* follows a concentration gradient. Highly concentrated formomarinobactins are necessary to completely inhibit the growth *P. aeruginosa*. Furthermore, in iron-rich environments, the siderophores with shorter acyl chains (F_A_-F_D_) were more active than the longer acyl chains. Similarly, in iron-poor media there was an inverted trend between the length of the acyl chain and the inhibition diameter between F_C_ to F_F_. This observation implies that for *P. aeruginosa* both the chemical structure (acyl chain length) and iron availability influence the siderophore’s growth inhibiting effect. The dependence on acyl chain length could reflect differences in membrane interaction, uptake efficiency, or diffusion.

## 4 Discussion

Amphiphilic siderophores have, until recently, been rarely described in *Pseudomonas*. Corrugatin, the first such siderophore identified nearly three decades ago, consists of an eight-amino-acid peptide linked to a short C8 fatty acid chain (Risse et al. 1998). Subsequently, ornicorrugatin and histicorrugatin were discovered (Matthijs et al. 2008; 2016), these differ by only one or two peptide residues, and we propose grouping them as the corrugatin family. These early examples occupy the hydrophilic end of the amphiphilic spectrum, due to their short fatty acid chains and longer peptide moieties. More recently, lactuchelin was reported as a series of seven-amino-acid cores acylated with C8-C16 chains (Chesneau et al. 2025). Biosynthetic gene clusters for lactuchelin were detected in the genomes of only a handful of *Pseudomonas* species (Chesneau et al. 2025).

Another suite of amphiphilic siderophores is here described for the grass isolate *P. graminis* LMG 21661^T^ which produces structural analogues of marinobactin previously described in *Marinobacter* sp. (Martinez et al. 2000; Martinez and Butler 2007), to cope with iron limitation. These analogues share the same 6-residue peptide backbone as marinobactin but differ by formylation rather than acetylation of N-hydroxyornithine. In addition, the *P. graminis* formomarinobactin carries shorter hydrocarbon chains (C10 to C14) compared with the C12-C18 chains characteristic of marinobactin.

*Marinobacter* species are widely distributed in diverse marine and saline terrestrial environments, including low-temperature hydrothermal areas (Handley and Lloyd 2013). Their production of marinobactin may establish a hydrophobicity gradient wherein longer, more hydrophobic siderophores remain near the cell surface, while shorter, more hydrophilic variants extend further into the environment (Martinez et al. 2003). In fact, amphiphilic siderophores represent nearly half of known marine siderophores and are thought to reduce siderophore diffusion in oceanic settings (Boiteau et al. 2016).

Formomarinobactin-producing *Pseudomonas* strains have been isolated from a variety of terrestrial and freshwater sources, including plants (grasses, rhizosphere, seeds, fruits), chemical contaminated soils, insects, cloud water, lakes, and river waters. A single strain was isolated from seawater. The shorter acyl chains of formomarinobactins may therefore reflect an evolutionary adaptation to more terrestrial or freshwater habitats, where reduced hydrophobicity and membrane interactions are likely advantageous. Lactuchelin, which is also produced by strains from terrestrial or freshwater habitats (Chesneau et al. 2025), has a Ser-rich peptide chain (βOHAsp-Ser-Ser-Ser-Orn-Ser-βOHAsp) that is therefore highly hydrophilic, more than the formo/marinobactin core, and this could compensate for longer C8-C16 tails to maintain motility/solubility in soil.

Phylogenomic mapping of the formomarinobactin siderophore gene cluster revealed that the ability to produce formomarinobactin is not unique to *P. graminis* but is characteristic of strains of the *P. lutea* clade of the *P. fluorescens* super clade. The *P. lutea* clade is a relatively small clade which consists currently of six described species, but phylogenomic analyses in this study revealed a yet uncovered diversity and identified an additional 18 putative species. Genome mining in combination with mass analysis showed that all these strains are predicted or were shown to produce formomarinobactin, except for *P. bohemica* LMG 30182^T^ isolated from bark beetle (Saati-Santamaría et al. 2018), which lacks the biosynthetic genes, but does have a putative formomarinobactin receptor. Phylogenetic analysis indicated that the receptor of *P. bohemica* likely resulted from the loss of the biosynthesis genes. This pattern aligns with the Black Queen Hypothesis (BQH), where costly public goods such as leaky diffusible siderophores are produced by “helpers” (producers) but exploitable by “beneficiaries” (non-producers or cheaters) that retain uptake machinery, driving adaptive gene loss for biosynthetic pathways while conserving receptors for efficient scavenging (Morris et al. 2012). In fact, *P. bohemica* seems to be a more efficient scavenger than the other genome-sequenced strains of the *P. lutea* clade, possessing the highest number of TonB-dependent receptors, with an especially high number of predicted siderophore TonB-dependent receptors.

*Pseudomonas* species typically produce one to three distinct siderophores and encode multiple TonB-dependent receptors that enable the utilization of xenosiderophores, siderophores produced by other bacterial or fungal species. BLAST analysis identified numerous *Pseudomonas* strains harboring a putative formomarinobactin receptor, revealing a substantial capacity within the *P. fluorescens* super clade to exploit formomarinobactins as an iron source. Growth stimulation by formomarinobactin was confirmed for select *Pseudomonas* strains, with functional validation in *P. rhodesiae*. In addition, formomarinobactin growth stimulation was demonstrated for a *Phytopseudomonas* strain. Further testing with the analogue marinobactin, along with mutational studies in selected receptor-expressing strains, would provide deeper insights into substrate specificity and physiological relevance.

This growth-promoting effect is consistent with the high iron-chelation capacity reported for marinobactin siderophores. Although marinobactin-E exhibits a higher intrinsic Fe(III) stability constant (log K_ML_ = 31.8, Zhang et al. 2009) than pyoverdines’ conditional stabilities at pH 7 (log K_pH7_ = 24.26-27.14, Ongena et al., 2002), protonation of the hydroxamate ligands reduces its effective affinity, making it roughly comparable (∼26.7) under physiological conditions. The predicted high affinity of formomarinobactins could explain their exclusive occurrence in pyoverdine-negative strains, as maintaining two high-affinity siderophore systems would impose excessive biosynthetic and regulatory costs on the cell. A similar pattern is observed with lactuchelin (Chesneau et al. 2025), implying that both siderophores act as primary iron-chelating systems in their native hosts. This is in contrast with siderophores of the corrugatin-family. Except for corrugatin, which is produced by a pyoverdine non producing species, all the reported cases are in pyoverdine producing *Pseudomonas* strains. These siderophores therefore generally coexist with pyoverdine and likely assume a secondary role rather than replacing pyoverdine as the principal Fe(III) scavenger.

Additionally, growing evidence indicates that siderophores can chelate other metal ions such as Cu, Ni, Zn, Ga, Al, etc. and some heavy metals (Arnold 2024; Schalk et al. 2011; Gomes et al. 2024). Hydroxamate-type ligands are known to allow, albeit with lesser affinity, binding with other tri-charged cations such as Ga^3+^ or Al^3+^ (Evers et al. 1989) as well as Zn^2+^, Ni^2+^ and Cu^2+^ and Co^2+^ (Hernlem et al. 1996). However, this binding does not equate efficient uptake by receptors. The biosynthetic gene cluster and receptor of formomarinobactins are not regulated by Zn or Ni but only Fe, suggesting they are true siderophores. Nonetheless, they may still be able to chelate other metal ions due to their ligand type. Several *P. lutea* strains isolated from metal-contaminated soils attest to their proliferation in the presence of toxic metals which could indicate siderophore-based remediation strategies (Arnold 2024; Schalk et al. 2011; Hesse et al. 2018; Gomes et al. 2024).

Moreover, the growth inhibiting activity of members of the *P. lutea* clade can largely be attributed to their siderophore production. Growth inhibition of *S. capitis* I, *B. zanthoxylo* RSN02 and *P. aeruginosa* LMG 1242^T^ by the *P. lutea* clade strains was exclusively observed in iron-limited conditions. In direct contact with the purified formomarinobactins, the growth inhibition was present in both iron-limited and iron-rich media, although inhibition zones were smaller in iron-rich conditions.

Similarly, the production of lactuchelin from *P. lactucae* CFBP 13502 was necessary for the growth inhibition of *Xanthomonas campestris* pv. *campestris* 8004 (Xcc8004). Co-culture of Xcc8004 with lactuchelin BGC deletion mutant of CFBP13502 and supplementation of the non-mutant co-culture supernatant with 0.39-50 µM FeCl_3_, tested independently, resulted in the total and gradual loss of inhibition, respectively (Chesneau et al. 2025). A gene encoding for the manganese exporter protein, *mntH*, in Xcc8004 was shown to have been significantly altered in the presence of *P. lactucae* in iron-limited conditions suggesting an impairment in the Mn/Fe regulation of Xcc8004, prompting authors to argue that iron starvation led to the observed antagonism.

Growth inhibition results for *Enterobacteriaceae* sp. DV 4951 highlighted the presence of other molecules or a siderophore-independent mechanism involved in the inhibition of the strain by *P. lutea* clade strains. The culture, cell-free supernatant and cell suspension of *P. graminis* CPA-7 were previously shown to have no antagonistic effect *in vitro* against *E. coli* O157:H7 or *Salmonella*, while CPA-7 dose-dependently reduced the log_10_(cfu) populations of both strains when inoculated on fresh cut apple and peaches (Alegre et al. 2013b). The authors suggested a competitive reduction rather than the secretion of antagonistic molecules. However, iron concentration was not considered and may explain the lack of activity of CPA-7 in medium against *E. coli* and *Salmonella* by the absence of siderophore production.

*P. bohemica* exclusively possesses anti-yeast activity contrary to other *P. lutea* clade strains. An iron regulated molecule is likely responsible for this activity because antifungal activity was iron-repressed. *P. bohemica* has previously been shown to inhibit the growth of *Candida humilis* (Saati-Santamaría et al. 2018). This selective activity emphasizes *P. bohemica*’s potential as a targeted antagonist against specific yeasts.

All in all, the iron chelating efficiency combined with the growth inhibiting properties of amphiphilic formomarinobactin produced by *P. graminis* and other strains of the *P. lutea* clade shows that siderophore production is not only a mechanism of iron acquisition but also a critical ecological trait that structures the competitive dominance and biocontrol potential of the *P. lutea* G, including *P. graminis*, especially, though not exclusively, in iron limited conditions.

## Author Contributions

CG, ML and KH performed the growth inhibition assays, purified and tested the siderophore, and carried out LC-MS analysis. BC and SM did the phylogenetic analysis. SM did the growth stimulation assays. ML and NB performed the RT-qPCR experiments and constructed the mutants. KH carried out LC-MS/MS analysis and elucidated the structure of formomarinobactin. Data analysis and manuscript write-up was done by KH and SM. SM conceived and supervised the entire study. All authors reviewed and proofread the manuscript.

## Acknowledgments

Undine Behrendt, Roeland Berendsen, Sigrid Flahaut, Rofida Benayad, and Ruben Werquin are acknowledged for the generous gift of strains. Pierre Van Antwerpen and Axelle Bourez for access to the Analytical Platform of the Faculty of Pharmacy of ULB and practical guidance.

