## Supplementary Figures for "Structures and broad-spectrum growth-inhibiting activity of formomarinobactin, formylated marinobactin analogues from the *Pseudomonas lutea* clade"

**Supplementary Table S1**. Primers used for the construction of the in-frame deletion mutants Prho$^{T}$ΔpvdLΔtrbABC and Prho$^{T}$ΔpvdLΔtrbABCΔTWR49273 of *P. rhodesiae* LMG 17764$^{T}$. Restriction sites are underlined.

| Primer name | Primer sequence (5'→3') |
|---|---|
| **Construction ΔpvdL:** | |
| Amplification of fragment A and B | |
| Prho$^{T}$-pvdL-AF | GTGAAGCTTCATGATGGACGCCTTCGAACT |
| Prho$^{T}$-pvdL-AR | GTGTCTAGAGAAGCGCTCCAAGGTGTCAA |
| Prho$^{T}$-pvdL-BF | GTGTCTAGAGCCTCGACCCGTGGCACTT |
| Prho$^{T}$-pvdL-BR | GTGGAATTCCCCTTCCAACTCCGCCATCA |
| Confirmation *pvdL* deletion mutant | |
| Prho$^{T}$-pvdL-conF | CGACGGTCACCACTTCTTCA |
| Prho$^{T}$-pvdL-conR | GCCATCGAGGCGTGGTATC |
| **Construction ΔtrbABC:** | |
| Amplification of fragment A and B | |
| Prho$^{T}$-trbABC-AF | GTGAAGCTTGGCAGCTGATCGATCGATAC |
| Prho$^{T}$-trbABC-AR | GTGTCTAGAAGGGACGATTCAGGCGTCAT |
| Prho$^{T}$-trbABC-BF | GTGTCTAGACTTGAGTTACGCCGTGGATTG |
| Prho$^{T}$-trbABC-BR | GTGGGATCCCCGATTTATCGAACAGGCATTG |
| Confirmation *trbABC* deletion mutant | |
| Prho$^{T}$-trbABC-conF | GGCTCCATCGACGCCATCA |
| Prho$^{T}$-trbABC-conR | GTGCGCGAAGAGTTGGATGA |
| **Construction ΔTWR49273:** | |
| Amplification of fragment A and B | |
| Prho$^{T}$-TWR49273-AF | GTGAAGCTTGGGCTATCTCGGTTGGAATC |
| Prho$^{T}$-TWR49273-AR | GTGTCTAGAGGATGGGGAGGCGCATTG |
| Prho$^{T}$-TWR49273-BF | GTGTCTAGATGCTCGGTACGGTGTCTTAC |
| Prho$^{T}$-TWR49273-BR | GTGGAATTCCGACGAGTTATCCACAGTTCT |
| Confirmation TWR49273 deletion mutant | |
| Prho$^{T}$-TWR49273-conF | TGCCTGGCAGCAATGGTTGA |
| Prho$^{T}$-TWR49273-conR | GTGCATAACCTTACGGAACAGTT |

**Supplementary Table S2.** NRPS adenylation domain specificity prediction using antiSMASH. X = unassigned residue

| Module | PKS/NRPS | Nearest Stachelhaus code | 8Å match | Stachelhaus code match |
|---|---|---|---|---|
| MrbD | | | | |
| 1 | D-Asp | DLTKIGHVGK | 88% | 100% (strong) |
| 2 | Dab | DIWELTADDK | 88% | 100% (strong) |
| MrbE | | | | |
| 1 | D-Ser | DVWHVSLIDK | 97% | 100% (strong) |
| 2 | X = Fo-OH-Orn | DGEVCGGVTK | 74% | 80% (moderate) |
| | X = OH-Orn | DGEACGGVTK | 71% | |
| 3 | Ser | DVWHVSLIDK | 97% | 100% (strong) |
| 4 | X = Fo-OH-Orn | DGEVCGGVTK | 74% | 80% (moderate) |
| | X = OH-Orn | DGEACGGVTK | 71% | |

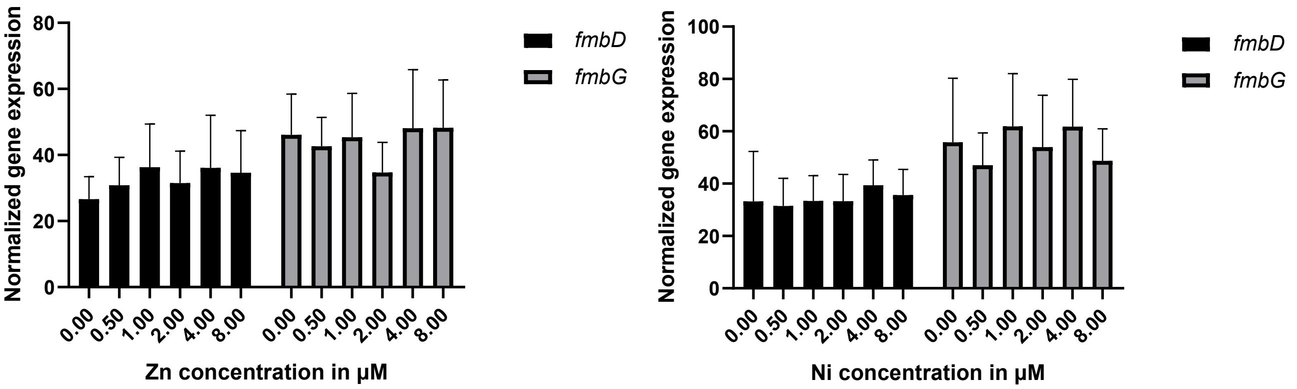

**Supplementary Figure S1.** Relative expression of *fmbD* and *fmbG* genes with increasing Zn (left) and Ni (right) concentrations normalized against reference gene *nadB*. Experiments were performed in triplicate. Values correspond to mean ± SEM. Brown-Forsythe and Welch ANOVA followed by Dunnett's T3 test for multiple comparisons indicates non-significant differences between control (0 µM) and higher concentrations.

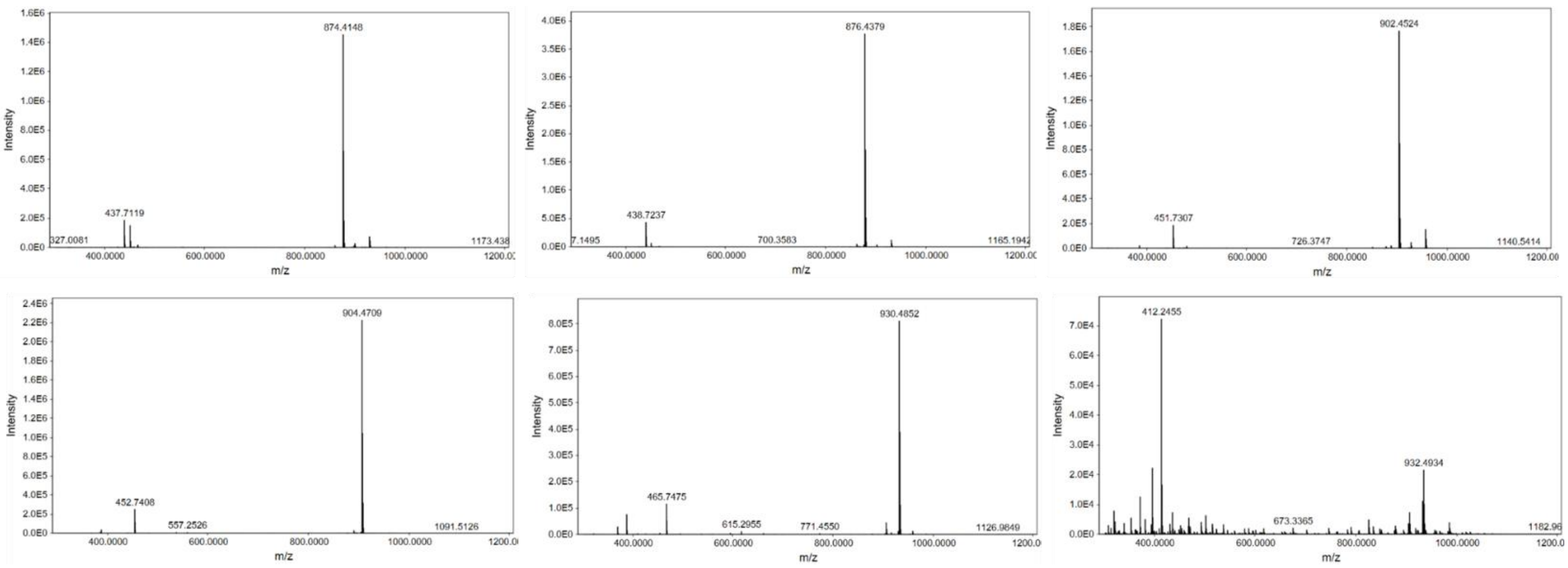

**Supplementary Figure S2.** Mass spectrum of $[M+H]^+$ and $[M+2H]^{2+}$ of each formomarinobactin peak *m/z* 874, 876, 902, 904, 930 and 932.

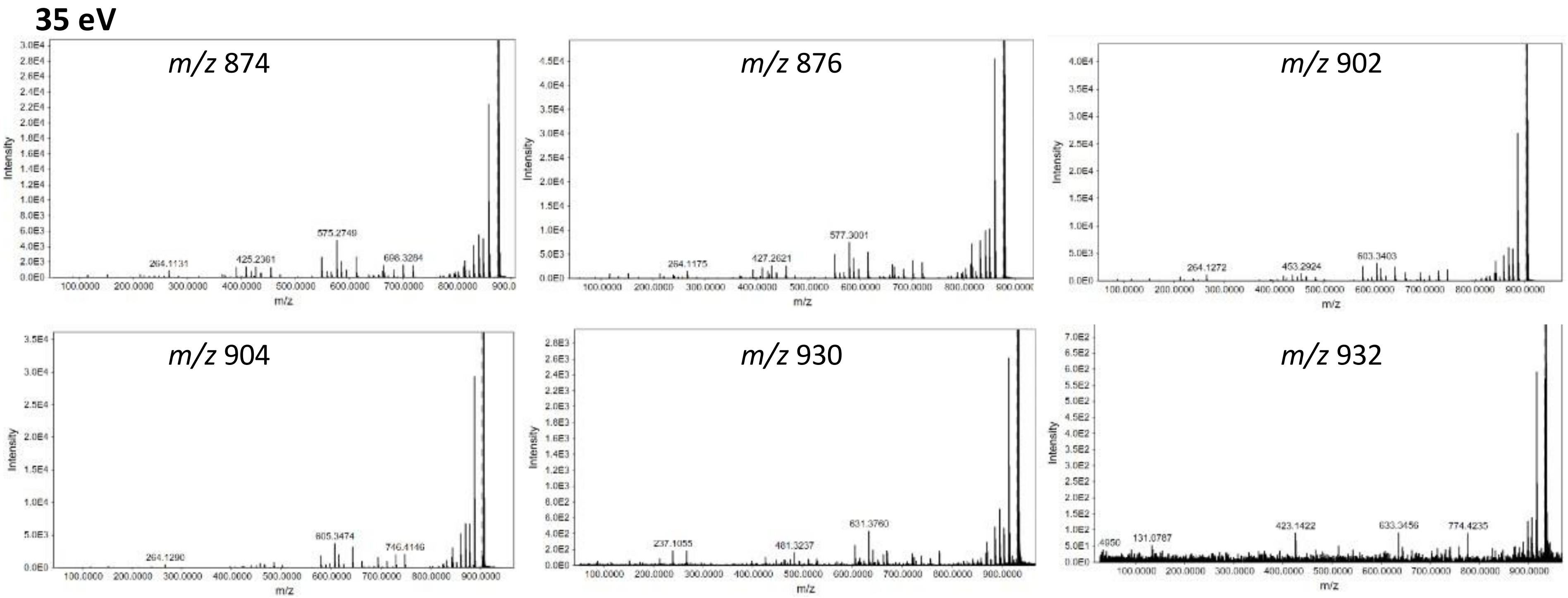

**Supplementary Figure S3a.** MS/MS Spectra of the formomarinobactins at CID 35 eV. Similar fragmentation patterns are observed.

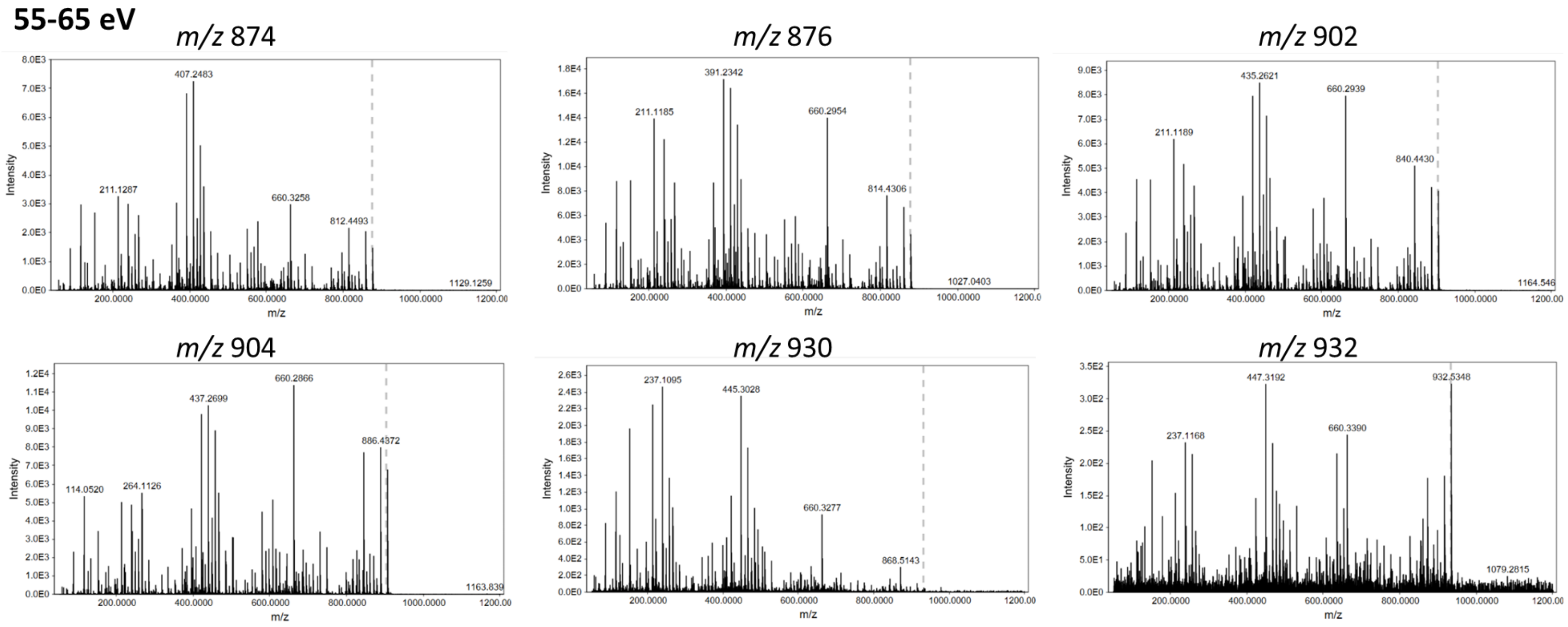

**Supplementary Figure S3b.** MS/MS Spectra of the formomarinobactins at CID 55-65 eV. Common fragment at 660 Da.

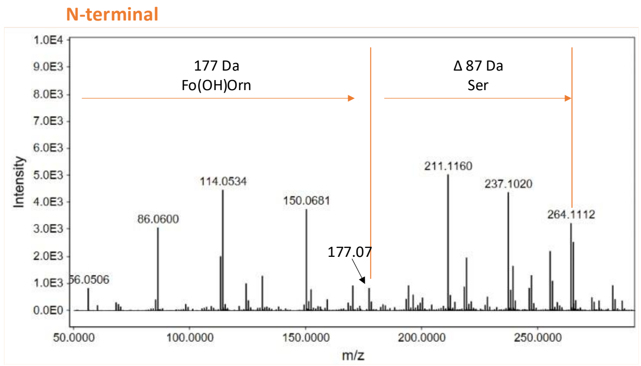

**Supplementary Figure S4.** Fragmentation of *m/z* 874 at 55 eV (CID) below 264 Da showing common peaks between formomarinobactin variants.

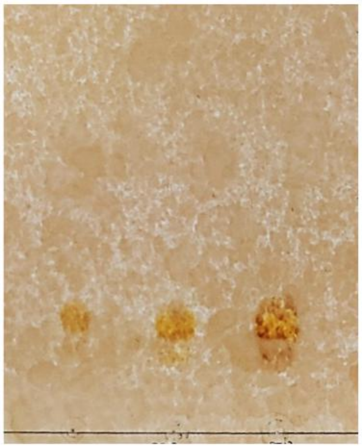

**Supplementary Figure S5.** Thin layer chromatography of semi-purified formomarinobactin. 5, 10 and 20 µl samples were spotted on precoated TLC silica gel 60 F254 plates (Merck) and were allowed to dry. The plates were run in an n-butanol:acetic acid:$dH_2O$ (12:3:5) solvent system**.**

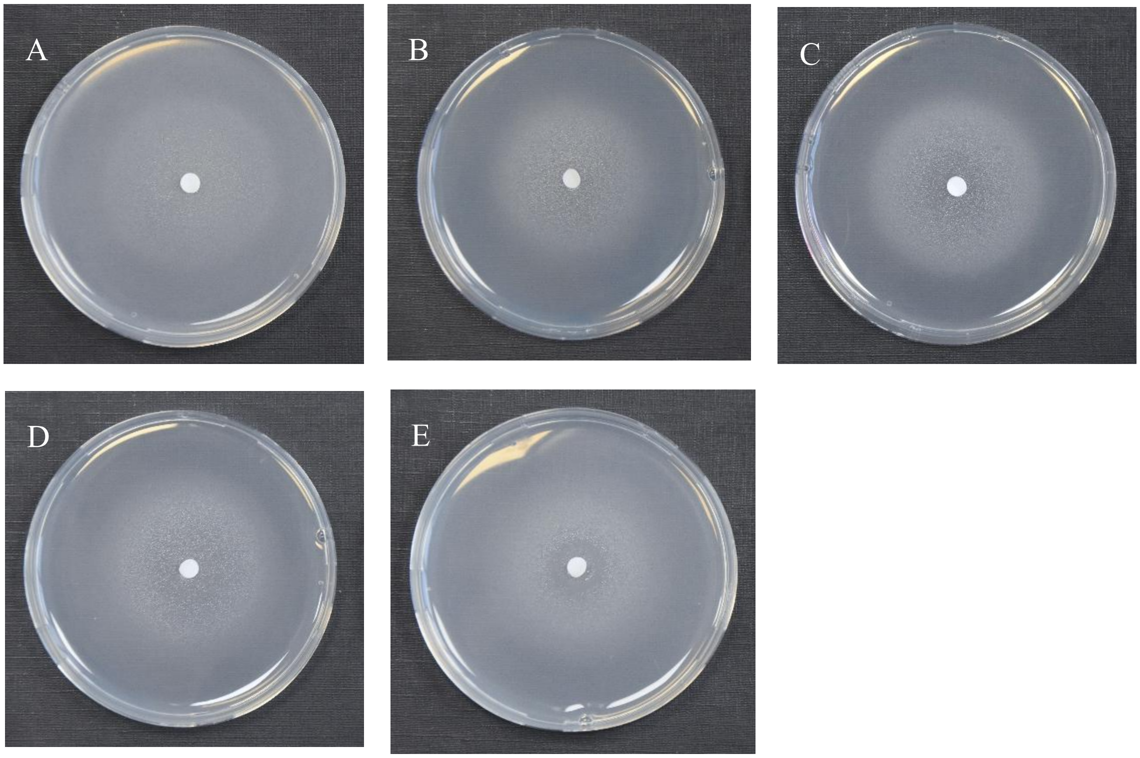

**Supplementary Figure S6.** Supplementary Figure S6. Growth stimulation assay of pyoverdine-negative *Pseudomonas* type strains and siderophore-negative *Pseudomonas* mutants on CAA medium supplemented with the iron chelator 2,2′-bipyridine. Pyoverdine-negative type strains: (**A**) *P. kuykendallii* LMG 26364$^{T}$ (200 µM). Siderophore-negative mutants: (**B**) *P. simiae* WCS417-M634 (500 µM), (**C**) *P. fluorescens* SBW25-15F3 (400 µM), (**D**) *Pseudomonas* sp. WCS374-BT1 (600 µM) and (**E**) *P. capeferrum* WCS358-JM213 (500 µM).

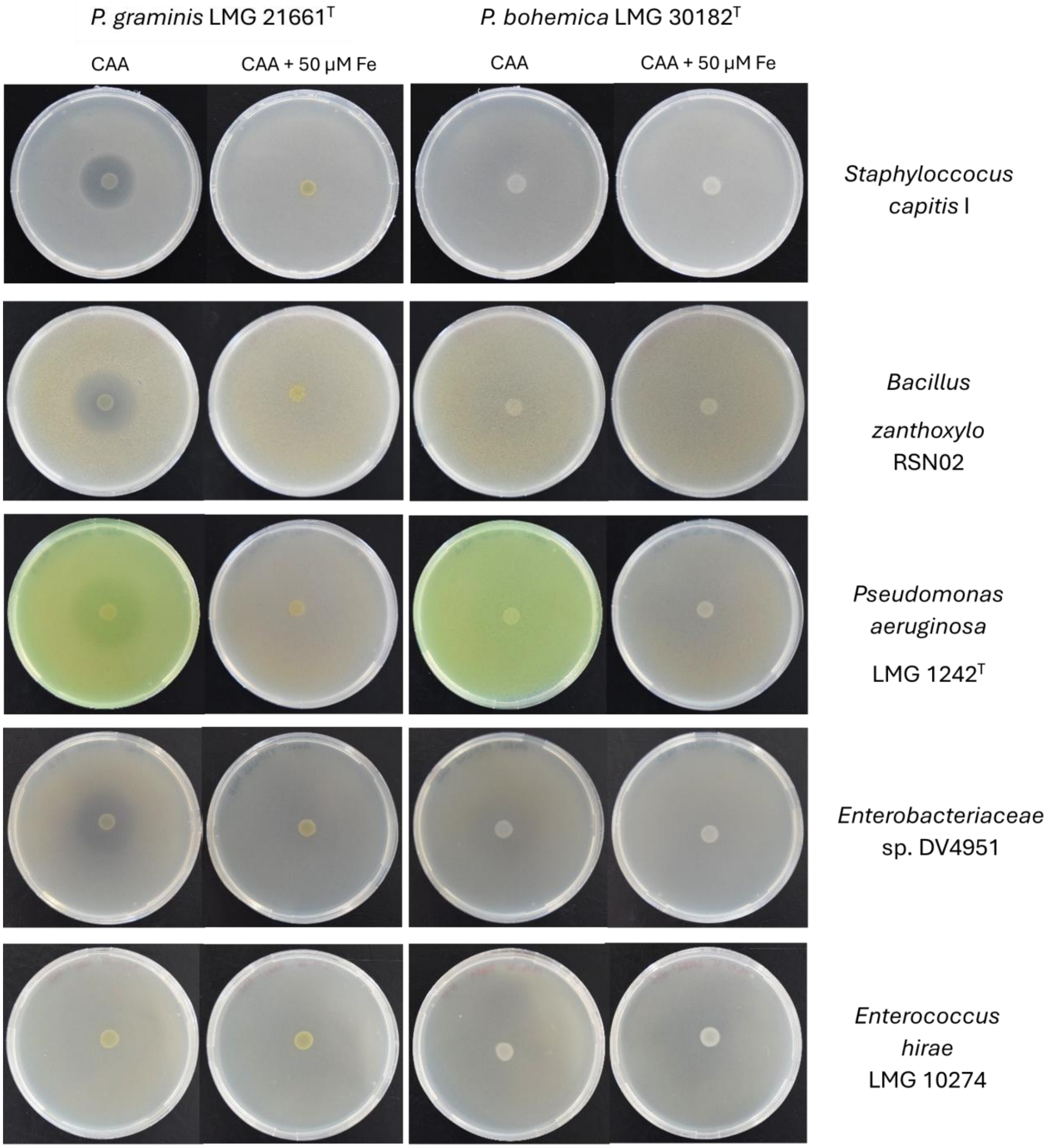

**Supplementary Figure S4.** Growth inhibition assay of *P. graminis* LMG 21661$^T$ (left column) and *P. bohemica* LMG 30182$^T$ (right column) against *S. capitis* I, *B. zanthoxylo* RSN02*, P. aeruginosa* LMG 1242$^T$, *Enterobacteriaceae* sp. DV4951 and *E. hirae* LMG 10274 (rows). All images are linearly adjusted for contrast (+20%) for presentation.

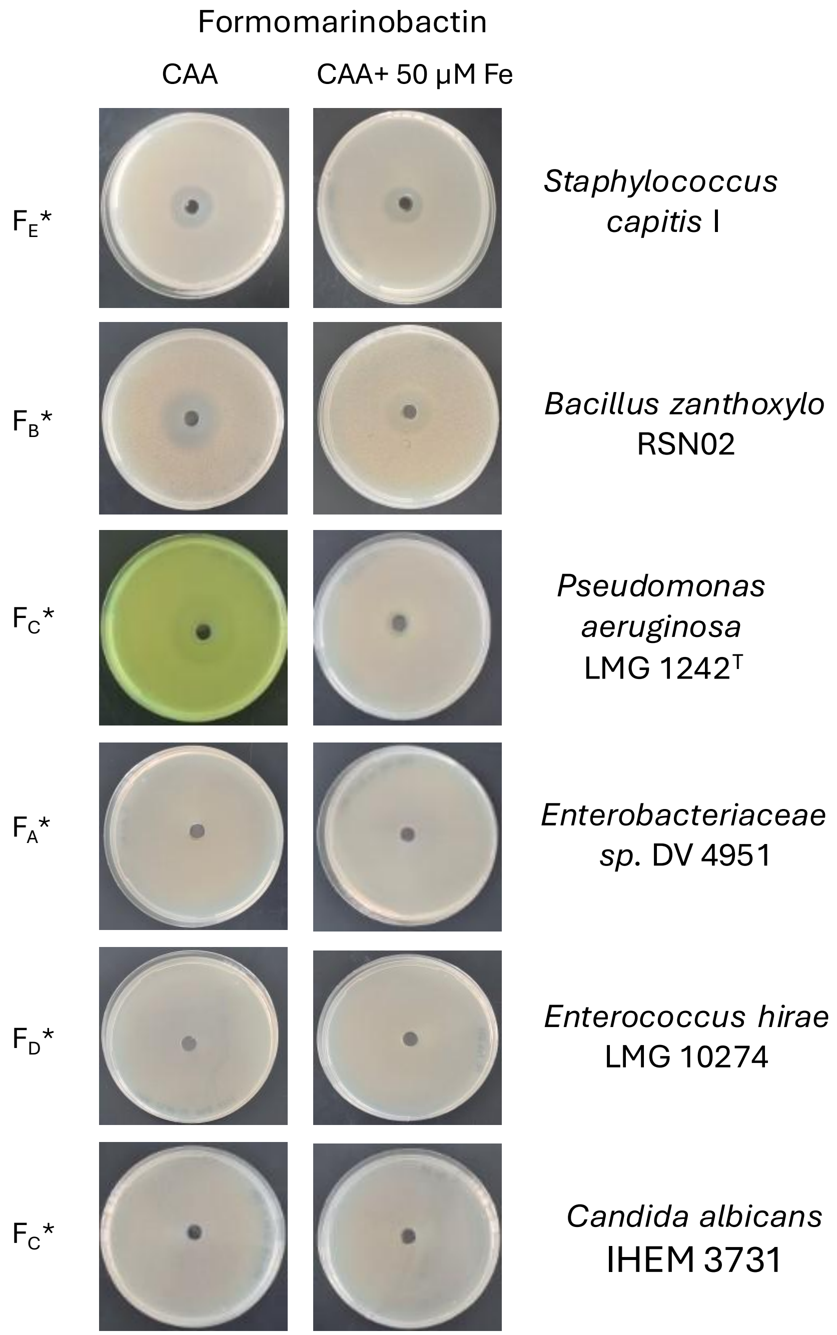

**Supplementary Figure S8.** Growth inhibition of formomarinobactins against *S. capitis* I, *B. zanthoxylo* RSN02, *P. aeruginosa* LMG 1242$^T$, *Enterobacteriaceae* sp. DV4951, *E. hirae* LMG 10274 and *C. albicans* IHEM 3731. *Molecule chosen for representative picture ($F_A$ = *m/z* 874, $F_B$ = *m/z* 876, $F_C$ = *m/z* 902, $F_D$ = *m/z* 904 and $F_E$ = *m/z* 930).
